# The Origin of the Ionic-strength Dependent Reentrant Behavior in Liquid-Liquid Phase Separation of Uncharged IDPs

**DOI:** 10.1101/2025.03.20.644249

**Authors:** Sayantan Mondal, Eugene Shakhnovich

**Author notes:** Corresponding author’s.

## Abstract

The effect of salt on coacervation of synthetic or biological polyelectrolytes and polyampholytes is well-studied. However, recent experiments showed that largely uncharged IDPs (like FUS) also undergo LLPS at physiological salt concentrations such as [C_ion_]∼0.15M, dissolve at higher salt concentration, and again phase separate at even higher salt concentrations such as, [C_ion_]∼3M. Here we use analytical theory and simulations to reveal the mechanism of these transitions. At low [C_ion_], the ionic solution acts as a highly correlated medium conferring long-range effective attractive interactions between spatially distant monomers. In this regime the ion concentration inside the condensate is higher than in the bulk solution. As [C_ion_] increases, the correlation length in the ionic plasma decreases, and the condensate dissolves. Second LLPS at high [C_ion_] is due to the entropy-driven crowding, and ion concentration inside the condensate is lower than in the bulk. Our study unravels a general physical mechanism of salt-dependent reentrant behavior in LLPS in uncharged IDPs.

## Introduction

Intrinsically disordered proteins (IDPs) and genetic materials can undergo liquid-liquid phase separation (LLPS) which leads to the formation of biomolecular condensates.^1^ The condensates exhibit liquid droplet-like behavior and underlie the formation of membrane-less organelles (such as nucleolus, stress granules, Cajal bodies, etc.). These organelles provide an additional means to compartmentalize subcellular processes.^2^ In addition to the intracellular compartmentalization, such condensates/droplets are involved in a variety of biological processes, such as genome reorganization and transcription,^3-5^ stress response,^6^ noise buffering,^7^ signal transduction,^8^ and membrane remodeling.^9, 10^ Disruptions in LLPS have been connected to the onset of numerous diseases, such as neurodegenerative conditions and cancer.^11^ During the past decade, there has been an increasing interest in understanding the formation, microscopic structure, and kinetics of LLPS, both experimentally and theoretically.^12-19^

Over the years, the molecular grammar of LLPS has been decoded.^20, 21^ Although new insights into the molecular grammar are emerging, it is well-accepted that individually weak but multivalent interactions between IDPs (such as, π − π stacking, cation-π, dipole-dipole, and charge-charge) are the major drivers of LLPS.^22, 23^ The condensation of IDPs are often favored by thermodynamics where the enthalpic gain through the formation of several weak multivalent interactions and entropic gain from the release of bound water/counterions surpasses the conformational entropy loss of the IDP and the de-mixing entropy loss.^24-26^ Although formation of a single (nearly) spherical phase is expected from a thermodynamic perspective, experiments always reveal a rather poly-dispersed multi droplet state that has been explained theoretically from a kinetics perspective.^17, 19^ The droplet size distribution was shown to obey a power law in some cases.^27^

The LLPS propensity of an IDP is governed by its sequence and patterning of charges including phosphorylation sites.^16, 18, 28-30^ Other than the sequence determinants, several physicochemical factors such as, temperature, presence of co-solutes, ionic strength, pH, etc. can alter the LLPS propensity. In some cases, an interesting phenomenon called ‘*reentrant phase separation*’ is observed, where the monotonic variation of a control parameter (say, ionic strength) transforms a system from a phase separated state to a macroscopically identical/similar phase separated state via two distinct transitions.^31^

The RNA-binding protein FUsed in Sarcoma (FUS) undergoes LLPS via homotypic interactions. FUS is found to be enriched in the nucleus and plays several important regulatory roles. The phase diagram of FUS condensation in the temperature-concentration space is known.^32^ In a recent study, Krainer *et al*. showed that FUS (along with four other IDPs) exhibits reentrant phase separation with respect to the salt concentration in solution.^33^ FUS and other IDPs show LLPS at low (physiologically relevant) as well as at high (physiologically irrelevant) salt concentrations but dissolve at intermediate salt concentrations. They suggested that, at low salt concentration regime, phase separation is driven by a mixture of hydrophobic and electrostatic interactions whereas, at high salt concentration regime, phase separation is driven by hydrophobic as well as enhanced non-ionic interactions. While their analysis provides a microscopic insight into the dominant interactions at various solvent conditions, the physical reason behind multiple reentrant transitions in a broad range of salt concentrations remain unclear. We notice that the five protein constructs used by Krainer *et al*.^33^ contain a significantly low *fraction of charged residues* (*f*_*+*_ and *f*_*−*_) and low net charge per residue (*NCPR*), especially in the disordered droplet-promoting regions. A detailed analysis based on their sequence is given in the **Supporting Information** (**Table S1**).

Previous studies highlighted the importance of salt in phase behavior of charged polymers, polyelectrolytes and polyampholytes, including IDPs.^34-38^ All these studies offered an intuitive and mechanistic explanation of the effect of salt on complexation (coacervation) of charged polymers: salt/ions screen attraction between monomers of opposite charges of the charged of a polyampholyte leading to de-mixing. In the same vein at higher salt concentration screening of this attractive interaction becomes more effective and reentrant transition into the dilute phase ensues. Apparently, counterions and salt ions in polyampholytes tend to weaken or altogether prevent de-mixing, which is most effective at no salt conditions in polyampholytes. However, this explanation is not applicable to mostly uncharged IDPs such as FUS and TDP-43 which also show salt-dependent reentrant LLPS.^33^ Thus, the mechanism of salt-dependent LLPS of an *uncharged* IDP remains unresolved.

Here we use explicit ion/solvent coarse-grained (CG) simulations combined with analytical statistical mechanical theory to unravel the physical origin of the reentrant LLPS in FUS and other uncharged IDPs. First, we present the observations from long-timescale coarse-grained simulations that explicitly take ions and water molecules into account and discern the essential mechanism(s) by running a series of control simulations where some factors (e.g. remaining FUS charges or salt ion charges) are abrogated one by one. Next, we present the detailed theoretical analysis based on the modified Voorn-Overbeek approximation^21^ to understand the simulation observations of FUS polymer in complex water salt solutions. Statistical mechanical analysis predicts two reentrant LLPS transitions in a range of salt concentrations observed in experiments/simulations as well as the distribution of ions inside and outside of the LLPS droplet in each LLPS regime. Overall, by using these two complementary approaches, we arrive at a general explanation of the reentrant LLPS phenomenon in uncharged IDPs. Although primarily motivated by the LLPS of FUS, the framework can be extended to other IDPs with a low number of charged amino acid residues.

## Materials and Methods

The conformational space of an IDP, such as FUS low-complexity domain (FUS-LCD, residues 1-163) is vast. Hence, molecular dynamics simulations at atomistic resolution are computationally expensive and far from probing the biologically relevant length- and timescales. Therefore, several coarse-graining strategies have emerged over the years. Here, we use the MARTINI v3.0^39^ general purpose coarse-grained (CG) force field to model our system and correlate with the results from the analytical theory. We choose the MARTINI model due to its recent success in understanding IDP dynamics and LLPS.^9, 40-45^ In addition, MARTINI has an explicit representation of the ions and water molecules (missing in other implicit solvent CG force fields) that are essential to the present study as discussed in the previous section. In the **Supporting Information** (section 5), we discuss the feasibility of simulating MARTINI at high salt concentrations by comparing it with an atomistic model that predicts NaCl solubility ∼ 6M. Below we outline the technical details and simulation protocols.

First, we create the structure from the sequence of human FUS-LCD, available in UniProt^46^ database (ID: P35637), by using PyMOL.^47^ Then the atomistic coordinates are converted into the MARTINI CG model with the help of the martinize2^48^ code. We randomly insert 42 copies of FUS-LCD chains inside a cubic box of dimension (30 nm)^3^. Following that we add the required number of counterions (2 Na_+_ for each FUS chain) and excess ions (Na_+_ and Cl_-_) to maintain electrostatic neutrality as well as to maintain the desired ionic strength in the range of 0.0 M to 3.0 M. We note that one MARTINI water bead is equivalent to 4 molecules of water. We note that the experiments used FUS concentration in the order of 10 μM. At this concentration, for one FUS chain we need an approximately (55 nm)^3^ large box and for the condensation simulations (say, with 25 chains) we need to simulate a box as large as (160 nm)^3^. As our model has explicit ions and water beads, such large systems will be computationally expensive, especially on the timescales we are interested in. Therefore, we used a concentration of FUS which has been previously simulated by Hummer *et al*.^*44*^ as well as Zerze *et al*.^45^ Although it is much higher than the experimental concentrations, it is not unrealistic in view of local enhancement of IDP concentration. As a result, the phase diagram/boundaries will change as well as the threshold value of salt concentration required for LLPS. In our study we find that the threshold lies in between 0.05 M and 0.10 M (**Supporting Information**, section 7). Therefore, the comparison with experiments is only qualitative.

As the experiments by Krainer *et al*.^*33*^ were performed using full length construct of FUS (526 residues), we additionally simulate LLPS of full-length FUS with the same MARTINI-3 parameters used to simulate FUS-LCD. Here we use 25 copies of full-length FUS inside a (50 nm)^3^ box filled with water and ion beads. During the simulation at three different salt concentrations, we preserve contacts in the folded regions (residues 285-371 & 422-453). We use the same simulation protocols as described in this section. The full-length FUS results agree well with the simulation observation of low-complexity FUS and are discussed in the **Supporting Information** section (Figure S6).

After a steepest descent energy minimization, we equilibrate each system for 10 ns followed by a 5 μs production run. We discard the first 2 μs from each trajectory and analyze the remaining 3 μs of the trajectory. The systems that exhibit LLPS formed a well-defined droplet within 1 μs whereas systems with no LLPS propensity remained dispersed throughout. Here we use slightly recalibrated force field parameters by increasing the protein-water Lennard-Jones interaction strength (ε_PW_) with a scaling factor λ = 1.03 that gives a reasonable transfer free-energy of FUS from the dilute to the dense phase, as also shown by Zerze *et al*.^45^

For equilibration and production run, we propagate the simulations with a timestep of 10 fs and 20 fs, respectively, using the leap-frog integrator. We use the modified Berendsen (V-rescale)^49^ thermostat (T=298 K and τ_T_=1 ps^-1^) and the Parrinello-Rahman^50^ barostat with isotropic pressure coupling (p=1 bar and τ_P_ = 12 ps^-1^) to maintain an NpT ensemble. For equilibration purposes, we use the Berendsen barostat^46^ with τ_P_ = 6 ps^-1^. The electrostatic interactions are screened with a dielectric constant (ε_r_) of 15 within and van der Waals interactions are terminated at 1.1 nm with the Verlet cut-off scheme. We perform all the simulations using GROMACS 2023.1 simulation package^51^ and conduct analyses with plumed 2.5.3.^52^ For visualizing the trajectories and creating snapshots, we use the visual molecular dynamics software (VMD 1.9.3).^53^

For the potential of mean force (PMF) calculations between two FUS monomers, we use well-tempered metadymaics,^54^ WT-MetaD (at T=300K and BiasFactor=5.0) with the inter center-of-mass distance (r_COM_) as the collective variable. We solvate two FUS chains in a 30 nm cubic box with MARTINI water and ions. We set the MetaD hill width to be 0.35 nm and the initial hill height to be 1.0 kJ/mol. The Gaussian hills are deposited every 1 ps and the MetaD simulations are run for 5 μs. We use PLUMED (v 2.9.0)^52^ patched with GROMACS,^51^ to perform the MetaD simulations and subsequent analyses. The PMF plots (**Figure 2**) were obtained by integrating the bias potentials using the ‘*sum_hills*’ utility in the PLUMED package.

**Figure 1.**
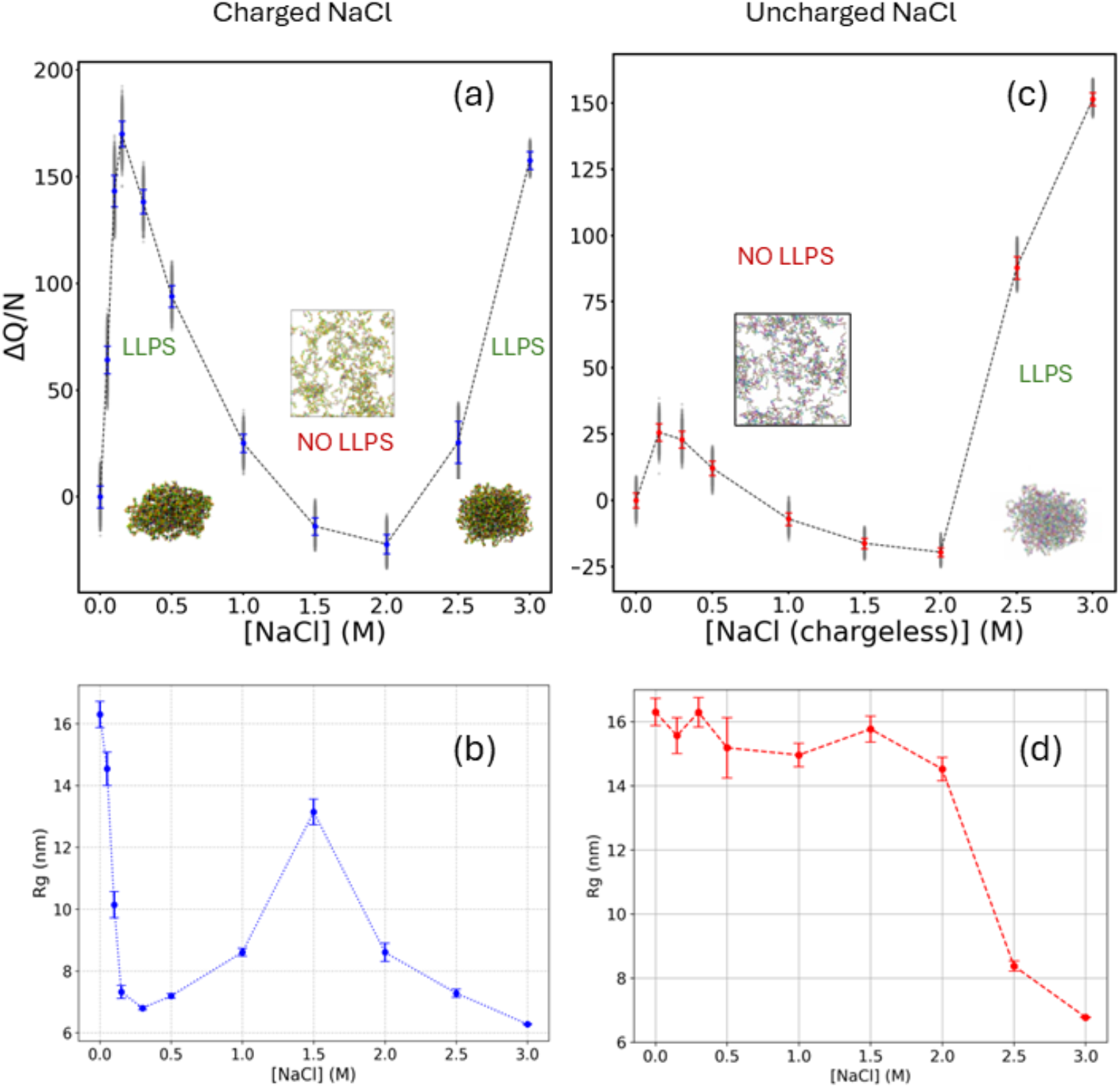
Time-averaged contact order parameter (ΔQ) per FUS chain against the ionic strength of the solution: (a) when sodium and chloride ions retain their charges and (c) when the sodium and chloride ions are made chargeless LJ beads. An increase in Δ)/N above a threshold indicates condensation. **Time averaged radius of gyration (for all IDP chains in the box), Rg against the ionic strength of the solution:** (b) when sodium and chloride ions retain their charges and (d) when the sodium and chloride ions are made chargeless LJ beads. A decrease in Rg indicates condensation. Both (a) and (b) panels show the ‘double reentrant’ phenomena. The dotted lines should be used as guides to the eye.

**Figure 2.**
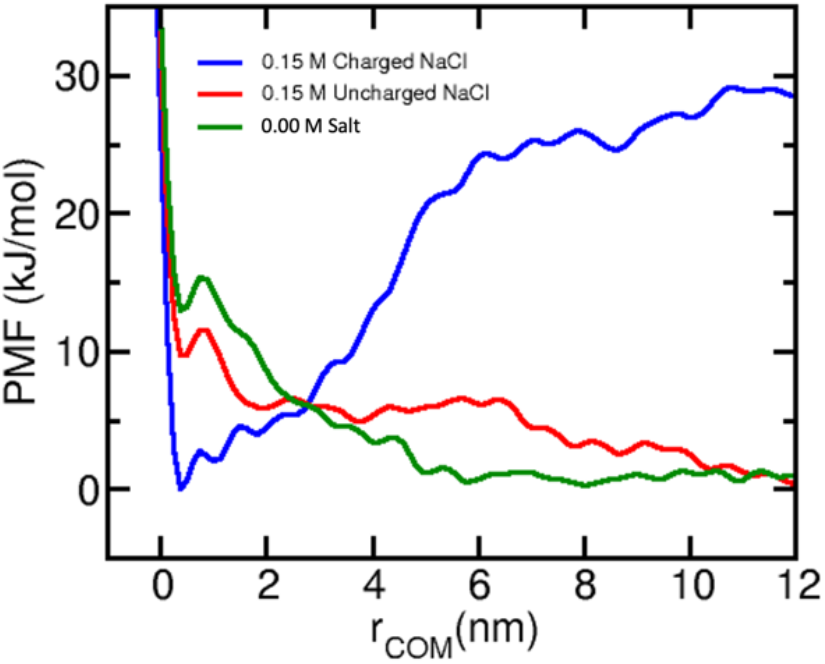
Effective attraction between FUS monomers through highly correlated medium of salt ions: Potential of mean force (PMF) between two FUS chains calculated by using well-tempered metadynamics. The two FUS chains are attractive in 0.15 M NaCl solution but becomes repulsive when the ionic charges are set to ‘0’, that is, electrostatics are turned off; or when no salt ions are present at 0.0 M NaCl.

Finally, to ascertain the reliability of MARTINI coarse-grained modeling, we perform atomistic simulations with CHARMM36 forcefield and TIP3P water model. We utilize the back-mapping algorithm to convert the equilibrated coarse-grained condensates into atomistic resolution. We have additionally modified the Lennard-Jones parameters of Na^+^ and Cl^-^ions following the work of Yagasaki *et al*., who obtained a solubility of 6.1 ± 0.30 mol/kg for NaCl in TIP3P water model.^55^ We started the atomistic simulations with a preformed droplet state and run the MD simulations up to 100 ns at 0.0 M, 0.15 M, and 3.0 M to check its stability. We note that starting from a dispersed phase would have been computationally challenging to simulate the timescales and length scales of interest. The results are presented in the **Supporting Information** (Section 8).

To quantify the extent of LLPS, we calculate a contact order parameter (COP), *Q*(*t*) as shown in **Eq. (1)**. The COP at a particular timeframe, *t* is expressed as:

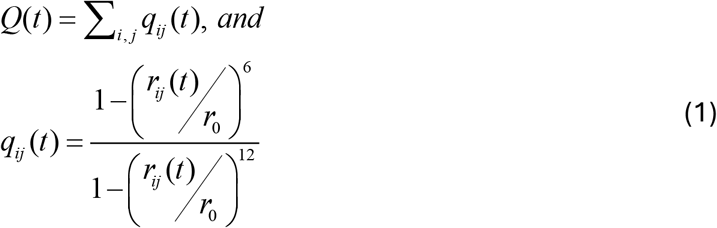

where *r*_*ij*_(*t*) is the distance between the i^th^ and j^th^ beads at time *t* and *r*_*0*_ is fixed to be 0.5 nm. Hence, *q*_*ij*_(*t*) can vary smoothly from 1 to 0 for a given pair. We further define Δ*Q* as the gain in COP by subtracting the time-averaged COP for FUS with only counterion system (**Figure 3d**), *Q*_0_. Therefore, Δ*Q* = ⟨*Q*(*t*)⟩ − *Q*_0_. The pair-interaction energies are obtained using ‘*gmx energy*’ code. The energies contain both Lennard-Jones and electrostatic interactions.

**Figure 3.**
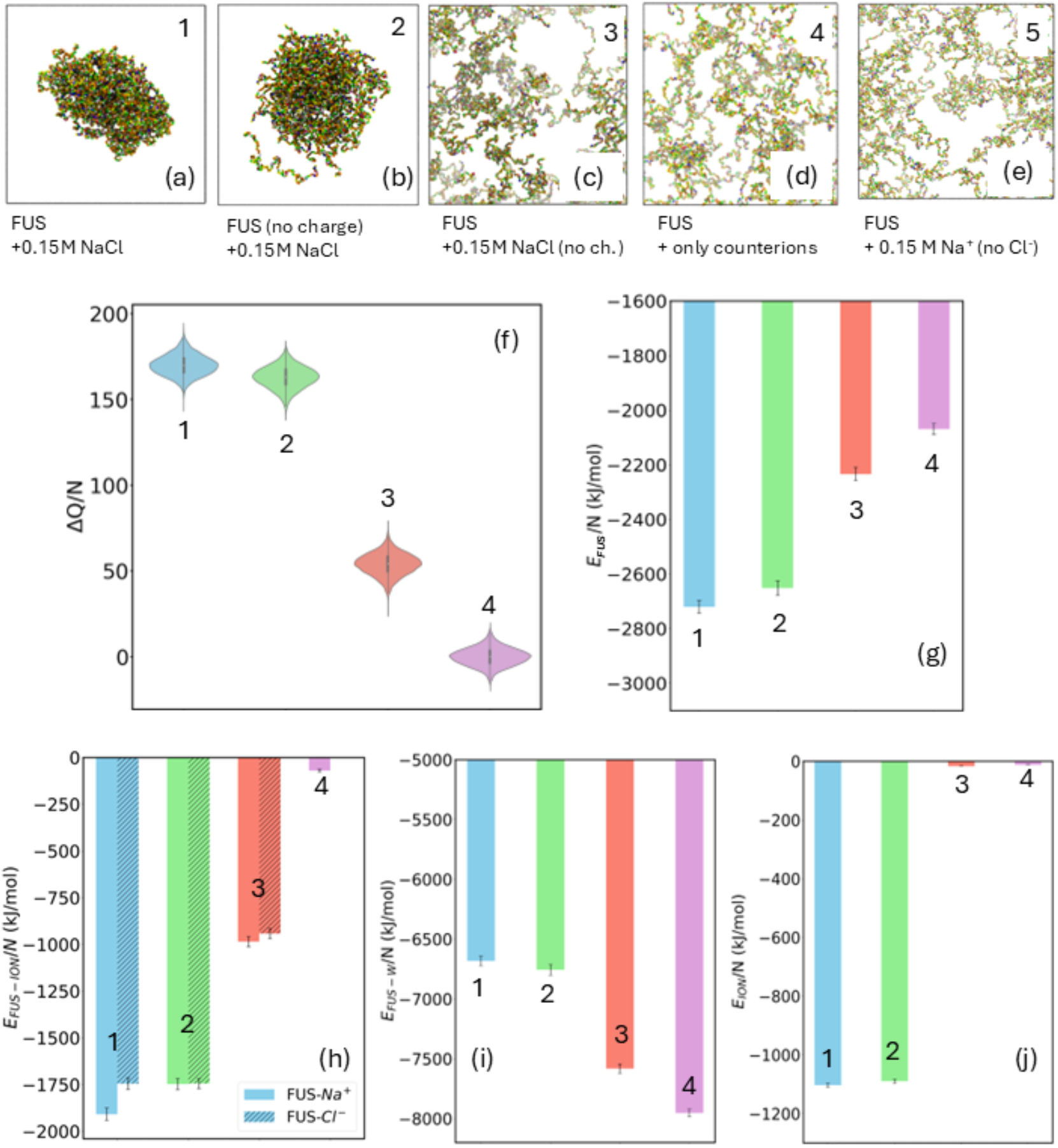
Representative snapshots and pair-interaction energetics from coarse-grained molecular dynamics: (a) FUS chains in 0.15M NaCl shows condensate formation, (b) Artificial FUS chains with no charges in in 0.15M NaCl shows condensate formation, (c) FUS chains in 0.15M artificial NaCl with no charges (which behave like LJ beads) shows no condensate formation, (d) FUS with only Na^+^ (2 ions per chain to maintain electrostatic neutrality) ions shows no condensate formation, and (e) FUS with 0.15 M excess Na^+^ (no Cl^-^) ions shows no condensate formation. The first four systems are respectively denoted as 1,2,3, and 4 in the subsequent plots. (f) Variation of the contact order parameter (Δ)) per FUS chain for the four systems. Time-averaged interaction energies of different pairs: (g) Among FUS chains, (h) Between FUS and ions, (i) Between FUS and water beads, and (j) Among the ions.

## Results

First, we study the propensity of FUS to undergo LLPS in the range of ion concentrations from 0.0 M to 3.0 M as observed in experiment.^33^ We observe no condensation at 0 M. Starting from physiological 0.15 M salt FUS undergoes LLPS into single compact cluster which dissolves beyond 1.0 M NaCl. Interestingly, LLPS reappears when salt concentration is above 2.5 M (**Figure 1a**). This ‘extremely high’ salt concentration regime, although biologically non-relevant, is important to understand from a physics perspective as noted below.

In **Figure 1a**, we plot the density of FUS Δ*Q*/*N* against salt concentration, [NaCl]. A sharp increase can be observed from [NaCl]=0.0 M to 0.15 M. The value of Δ*Q*/*N* decreases upon further increase of salt concentration and the LLPS cluster fully dissolves when [NaCl] exceeds 1.0 M. However, when [NaCl] > 2.0 M the value of Δ*Q*/*N* increases again and reaches the values observed at 0.15 M [NaCl]. This is the second entrance into the ‘LLPS’ regime from the ‘no LLPS’ zone. Therefore, we observe the double reentrant transition where the second reentrance is observed as in experiments.^33^ From the error bars (or, standard deviations) **Figure 1a**, important information regarding the dynamics of the condensates can be gleaned. Although condensation/LLPS occurs at around [NaCl]=0.15 M and at around [NaCl]=3.0 M, the condensate is more dynamic for [NaCl]=0.15 M system (more dispersed data) whereas, at [NaCl]=3.0 M the condensate is less fluctuating (less dispersed data). The same can also be gleaned from another order parameter, radius of gyration (Rg) of all the FUS chains in the box, shown in **Figure 1b**. Rg sharply decreases for [NaCl]∼0.00M-0.50M followed by an increase around [NaCl]∼1.0 M and again decrease at higher [NaCl].

In **Figure 1c** we plot the same quantity as in **Figure 1a** (that is, Δ*Q*/*N* vs [NaCl]), however, for an artificial control system with uncharged salt (essentially LJ beads). This is equivalent to ‘turning off’ the electrostatic interactions in the system. There is a slight increase in Δ*Q*/*N* at lower salt concentrations (probably due to depletion forces), but without condensation. The absence of condensation manifest itself in the high Rg values at that salt concentration (**Figure 1d**) and visual inspections confirm it. Interestingly, FUS undergoes LLPS at high salt concentration even without charged slat ions (effectively, LJ beads), indicated by a sharp increase in Δ*Q*/*N* (**Figure 1c**) and a decrease in Rg (**Figure 1d**). We note that these are Rg of all the proteins taken together and do not refer to a single chain Rg. The analytical theory, discussed later in the manuscript, also predicted that the system would undergo phase separation at high salt concentrations when electrostatic interactions are ‘turned off’ (**Figure 5e**). Plots for pairwise interaction energies against salt concentrations, namely *E*_*FUS*_, *E*_*FUS* − *Ion*_, *E*_*FUS − W*_, and *E*_*Ion*_ can be found in **Supporting Information** (**Figures S2** and **S3**).

These findings indicate that, paradoxically, salt ions are crucial for condensation of mostly uncharged FUS at physiological salt concentrations. To get further insight into the origin of this surprising effect we carried out the well-tempered metadynamics (WT-MetaD) calculations to obtain the potential of mean force (PMF) between two FUS-LCD chains in 0.15 M NaCl solution, with and without the ionic charges (see **Methods** for technical details). The resultant PMFs are shown in **Figure 2** where the blue trace shows an effective attraction between two FUS chains when electrostatics are ‘on’, with approximately 30 kJ/mol free energy stabilization in the associated state. In the same figure, the red and green traces show an effective repulsion between the FUS chains when electrostatics are either ‘off’ or at [NaCl]=0.0 M. This demonstrates the long-range (∼ 10 nm) effective attraction through the ionic media which is absent in the presence of inert crowders or no salt ions.

Next, we explore in detail the mechanism of LLPS of FUS including, crucially, the role of ions. For control and comparison, we study several model systems with the same box size and number of FUS molecules. The five systems are as follows: (1) Original model of FUS at 0.15 M NaCl, (2) artificial ‘chargeless’ FUS chains in 0.15 M NaCl where a few charges of FUS chains are abrogated, (3) FUS with artificial ‘chargeless’ 0.15 M NaCl whereby charges of salt ions are set to zero, (4) FUS with only few Na^+^ ions serving as counterions to FUS charges, and (5) FUS with only 0.15 M Na_+_ ions without any Cl_-_ions. The last system contains high amount of net positive charge. Below we detail the rationale behing studying these systems. The purpose of the control system-2 is to check whether the 4 charges on FUS (N-ter, C-ter, Asp-5, and Asp-46) drive LLPS in any way. It turns out that FUS, whose own charges are set to 0 forms condensate in the regular NaCl solution whose salt ion charges are intact (**Figure 3b**). We confirmed this observations for system-2 by starting from a completely dispersed initial state as well (**Supporting Information, Figure S4**). Therefore, we can rule out the role of FUS counterion mediated inter-polymeric interaction in the FUS LLPS. The purpose of system-3 is to test whether the ionic charges are required for LLPS in the low [NaCl] regime as predcited by the analytical theory (see below). In agreement with the theoretical predcition, the condensate does not form in this case (**Figure 3c**). The purpose of the control system 4 (**Figure 3d**) is to demonstrate that FUS, with only its counterions (that is, 2 Na^+^ ions per chain) cannot exhibit LLPS. Importantly in control system 5 (**Figure 3e**), where Na_+_ ions retained their positive charges and the Cl anions were absent, the highly correlated Debye-Huckel plasma does not exist and FUS did not form condensate. This supports the conclusion from the analytical theory (discussed below) that LLPS at low salt concentration is driven by the interactions through highly correlated ionic plasma. All in all we conclude that highly correlated system of oppositely charged ions, a Debye-Huckel plasma, is driving reentrant condensation and dissolution of FUS at low [NaCl]. This mechanism has significant similarity to polymer-salt-induced condensation of DNA or psi(ψ)-condensation,^56, 57^ discussed subsequently.

To quantify the extent of LLPS, in **Figure 3f** we plot Δ*Q* normalized by the number of FUS chains for the four systems depicted in **Figure 3a** through **Figure 3d**. For systems 1 and 2, the gain in number of contact pairs is at the maximum. On the contrary, for systems 3 and 4, the value of Δ*Q*/*N* is much lower. Note that these are not the absolute values mentioned in the **Methods** section. In **Figure 3g** we show the time-averaged pair interaction energies (LJ + Coulomb) between FUS chains, *E*_*FUS*_/*N*. As one can presume, the interaction between FUS chains is the strongest for system 1, and almost comparable for system 2. However, these energies are substantially lower for systems 3 and 4, where the interaction primarily arises due to intra-chain contacts. In **Figure 3h**, the energies of FUS-ion interaction are plotted, separately for Na^+^ and Cl^−^. The values for systems 1 and 2 are lower which indicate enhanced interaction of FUS with salt ions. The FUS chains of system 1 interact slightly stronger with the Na^+^ than with the Cl^−^ owing to the presence of two negatively charged aspartate residues in each chain. Such distinction is not present in system 2 where FUS is made chargeless. In system 3, the excess ions are chargeless but the counterions (2 Na^+^ per chain) are not. Hence, it shows enhanced interaction with Na^+^, but the values are almost 50% reduced. System-4 has no Cl^−^ and only Na^+^ counterions. A reverse trend is observed in **Figure 3i** where the energy of FUS with water are plotted. Systems 1 and 2 interact less with water compared to system 3 and 4. **Figure 3j** shows the interaction energies of ions. Here we find that ions interact favorably when LLPS occurs. Overall, from the pair-energetics we find that LLPS of FUS stabilizes the inter-FUS, FUS-ion, and inter-ionic interactions; and destabilizes the FUS-water interactions.

Next, we study the spatial distribution of ions. Observations from our CG simulations are presented in **Figure 4**. We find that, in LLPS occurring at low salt concentration the condensate will absorb more ions, and in the second LLPS occurring at higher concentration the condensate will expel ions. The analytical theory presented below, also predicts the same (**Figure 5b**-**Figure 5d**). For the condensate at [NaCl]=0.15 M, there is a significantly increased population of both Na^+^ and Cl^−^ ions in the interior of the FUS condensate, as seen from the representative simulation snapshots (**Figure 4a, Figure 4b**, and **Figure 4c**). In **Figure 4d, Figure 4e**, and **Figure 4f** we respectively plot the spatial density profile of FUS, sodium, and chloride along the X-dimension of the simulation box. The density profile clearly shows the excessive adsorption of ions by the FUS condensates. In case of full length TDP-43, Ingolfsson *et al*. found similar increased ionic concentration inside the condensate, from MARTINI-3 CG simulations.^58^

**Figure 4.**
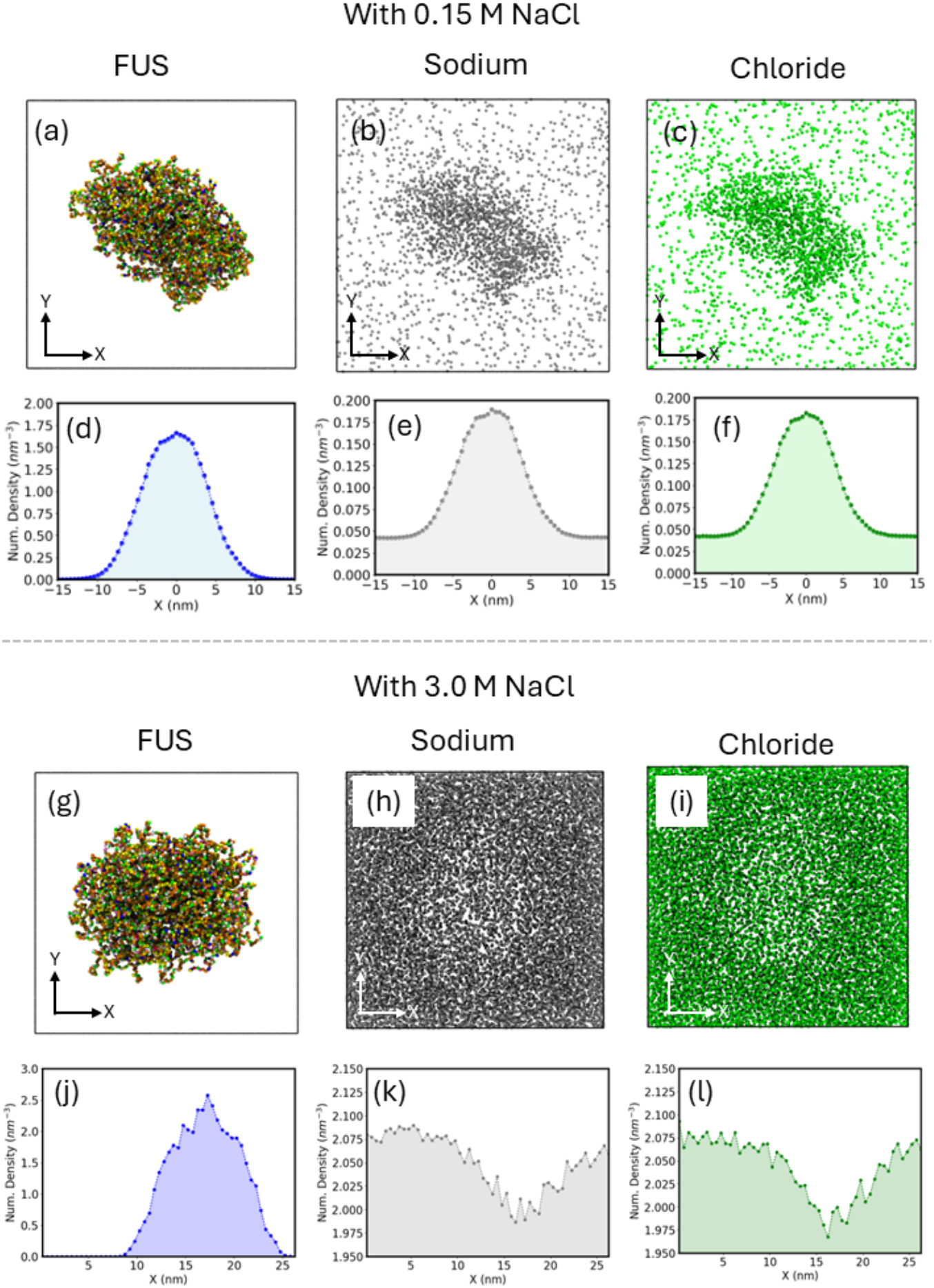
Spatial distribution of the FUS condensate, sodium, and chloride ions inside the simulation box for two different ionic strengths, namely, 0.15 M and 3.0 M. In the low ion concentration regime, the density of ions follows the density of FUS as shown pictorially in panels (a) – (c) and quantitatively from the spatial density profiles in panels (d) - (e). On the contrary, in the high ionic concentration regime, the FUS condensate expels ions from its interior as shown pictorially in panels (g) – (i) and quantitatively from the spatial density profiles in panels (j) – (l).

**Figure 5.**
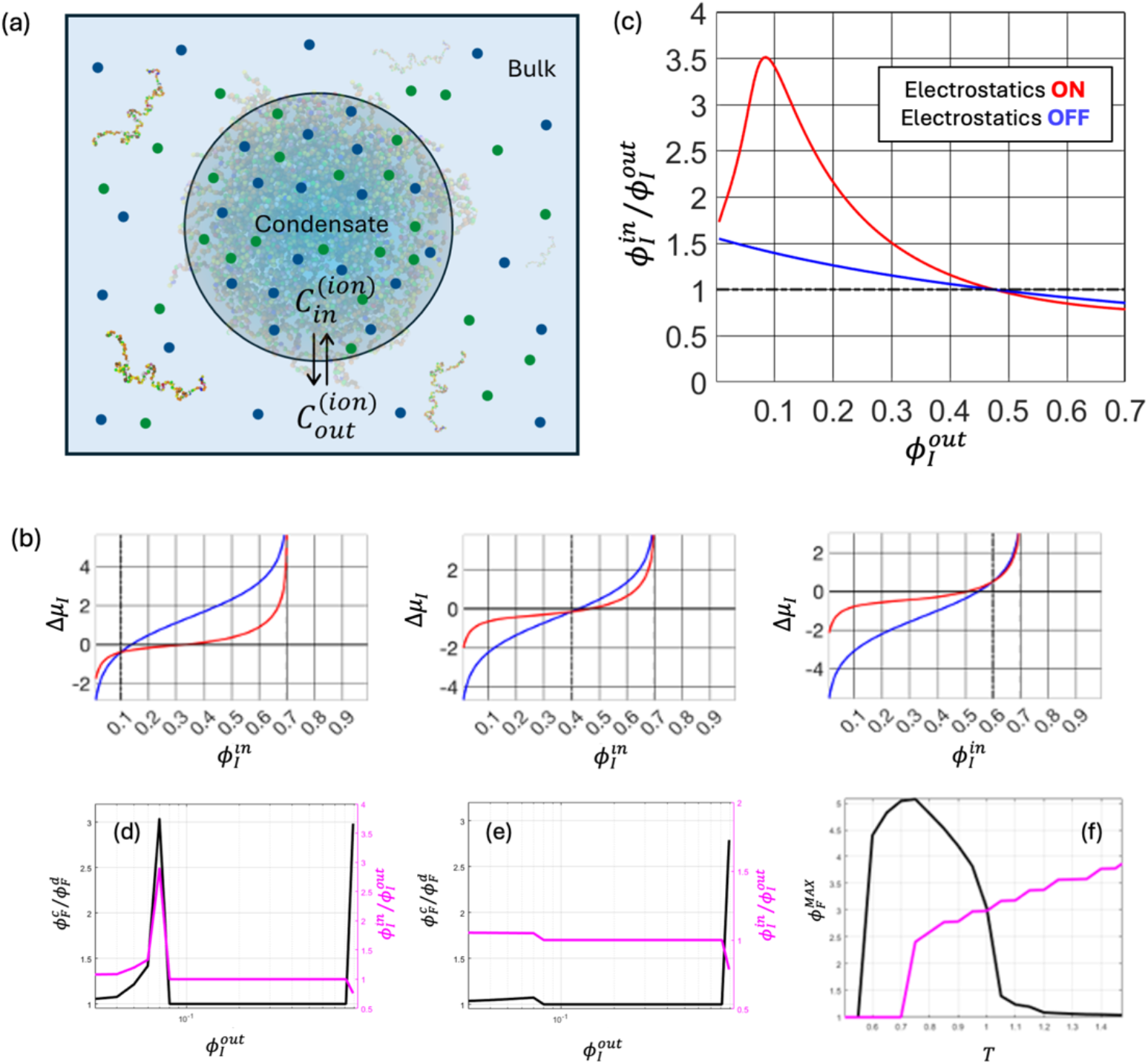
Statistical mechanics of liquid-liquid phase separation of uncharged polymers in water salt solutions: **(a)** A schematic illustration of the liquid-liquid phase separated system. The IDP rich phase is the condensate with volume fraction of ions 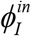 and the dilute/bulk phase with volume fraction 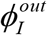. **(b)** Graphic solution of Eq.(3) for the difference of chemical potential of ions in the condensed phase vs. the bulk as a function of volume fraction of salt inside the condensate 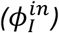. Three panels correspond to different values of 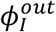 as marked by vertical dashed lines (0.1, 0.4, and 0.c respectively) Red lines correspond to full free energy functional with electrostatics between ions while the blue lines is a control where charges are “turned off”, i.e. A=0. The values of 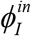 at which the lines intersect y=0 correspond to the solutions of Eq.(3) for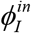. **(c)** Relative volume fraction of salt inside the condensate vs that in the dilute phase representing solution of Eq.(5) in full range of volume fractions of salt. **(d)** Reentrant transitions in the solution of FUS with 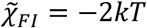. At low salt concentration FUS first condenses and as salt concentration increases condensates dissolve back into a one phase solution. Enrichment of salt inside the condensate drives the first transition until second phase separation transition accompanied by depletion of salt in the condensate. The Black curve shows the volume fraction of FUS in the condensate relative to dilute phase. The Magenta curve shows the volume fraction of ions in the condensate relative to the dilute phase. **(e)** same as **(d)** but with ion charges off (A=0). **(f)** Temperature dependence of the volume fraction in the FUS condensate in the low salt phase (black lines) and the high salt phase (magenta).

In the high salt concentration LLPS regime, for [NaCl]=3.0 M, the opposite effect is observed where the FUS-rich region of the box has lower density of ions than the region outside the condensate. It can be seen in the snapshots (**Figure 4g, Figure 4h**, and **Figure 4i**) and also from the density profile plots (**Figure 4j, Figure 4k**, and **Figure 4l**). This shows moderate expulsion of ions from the interior after reentrance. Also note that excessive adsorption of ions in the low salt regime is much stronger than their expulsion in the high salt regime (almost 3-fold for adsorption and ∼15% for expulsion), again in agreement with the predictions of the analytical theory as highlighted in **Figure 5** below.

### Theoretical Analysis

In this section we develop a simple mean filed theory to get a deeper insight into the findings from simulations that point out to the crucial role of NaCl ions in *promoting* LLPS in uncharged FUS-LC at low ionic strengths. These findings are somewhat surprising as it contradicts common intuition borne out of studies of net neutral polyampholytes and coacervating polyelectrolytes where salt ions weaken LLPS by screening the attraction between oppositely charged residues.^59^

Consider the phase-separated solution of an uncharged IDP (for example, LCD of FUS) which we model as an uncharged homopolymer. The dense phase occupies a volume *V* (together with water and ions) and the remaining volume (*V*_0_ − *VQ* is occupied by the dilute phase. We use the Voorn−Overbeek (VO)^35^ approximation to describe thermodynamics of uncharged polymer in the Flory-Huggins approximation and charged salt ions in the Debye-Huckel approximation. In addition to the usual electrostatic interactions in the VO theory we included direct *non-electrostatic* LJ interactions between uncharged FUS and salt ions. The total free energy of the system is given by:

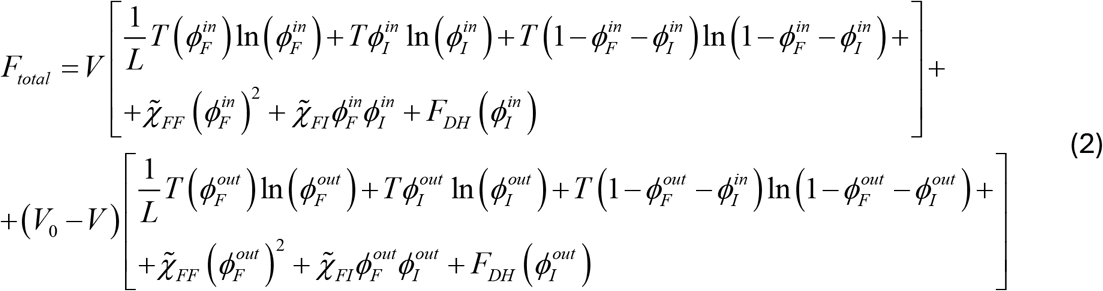

Where ϕ denotes volume fraction of ions and FUS inside the condensate or outside as indicated by corresponding subscripts/superscripts (see Supplementary Text and **Eq. S1** for definition of volume fractions). Square brackets denote free energy of FUS and ions in and out of the condensate. Flory-Huggins parameters 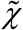 describe effective *non-electrostatic* LJ interactions between FUS, and the ions where explicit water is excluded via the incompressibility conditions given further discussed in Supplementary text.

We assume, for simplicity that salt anions and cations have equal non-electrostatic interactions with both water and FUS and do not make distinction between them. Electrostatic interactions are between salt ions only and they are described by the free energy contribution *F*_*DH*_ in the Debye-Huckel approximation as detailed below.

The ion-equilibrium between the condensate and the bulk is achieved at equal chemical potentials of ions in the condensate (“in”) and the bulk (“out”):

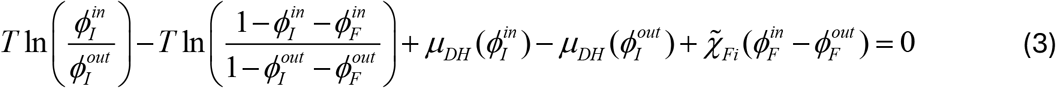

The DH chemical potential μ_*DH*_ is :^60^

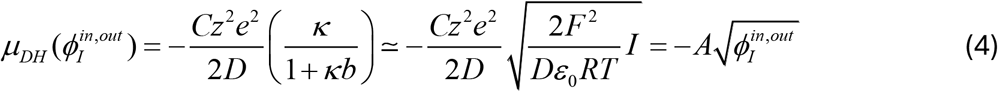

In **Eq.(4)**, *I* is the ionic strength of the salt solution defined as 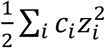 where *c*_*i*_ and *z*_*i*_ are the concentration and valency of the *i*^*th*^ ionic species, D is the dielectric permittivity of water, F is Faraday’s constant (= *eN*_*A*_), b is effective ion size, ε _0_ is the vacuum permittivity, and κ is the Debye length. At low to moderate salt concentrations κ*b* ≪ 1 and we disregard this term for the qualitative analysis below.

Equivalently the condition of equilibrium between FUS molecules in the bulk and condensate is that chemical potential of FUS molecules in the condensate is equal to that in the bulk 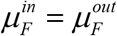 which brings in the equation that determines the volume fraction of FUS in the condensate at a fixed volume fraction of FUS in the bulk:

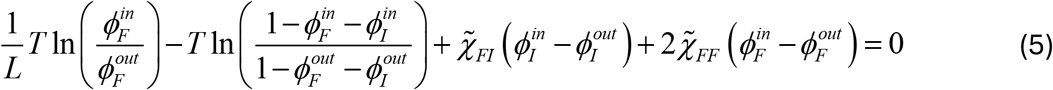

Where volume fractions of ions in the bulk and the condensate are determined by the equilibrium condition **Eq. (3)**.

Now we explore the whole range of densities for all components. To that end, in **Figure 5b** we plot **Eq. (3)** for the difference of chemical potentials, Δ*μ*_*I*_ *of ions* between condensed and dilute phases as a function of volume fraction of ions inside the condensate 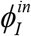 at various values of volume fractions of ions outside the condensate 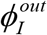. To illustrate the qualitative behavior of the chemical potential plots we fix the volume fraction of FUS in the dilute phase, 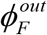 at 0.1 and inside the condensate, 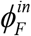 at 0.3 (other values give qualitatively similar results).

We see from **Figure 5b** that when salt concentration in the bulk is low, the condensate strongly absorbs ions and the effect is largely due to the Debye-Huckel correlation as cancellation of the DH energy in the free energy functional, **Eq.(2)** greatly diminishes absorption of ions in the condensate (blue curve). However, as the bulk salt concentration increases, this effect diminishes and finally reverses whereby the ions are depleted inside FUS condensate compared to the bulk. This can be summarized in **Figure 5c** that shows the dependence of ion concentration inside FUS condensate on the ion concentration in the bulk at a fixed density of FUS condensate. We note the non-monotonic dependence of concentration of the ions with full charge interactions inside the condensate (red line). In contrast, in the absence of the charge induced DH contribution to free energy the non-monotonic dependence disappears. Rather a much weaker monotonic dependence is observed (blue line in **Figure 5c**). For further analysis of the ion equilibrium including the limits on the reentrant behavior see the **Supporting Information** text.

Finally, we solve **Eq.(3)** and **Eq.(5)** at a fixed volume fraction of FUS in the dilute phase and obtain equilibrium density (volume fraction) of FUS in the condensate in a broad range of bulk ion concentrations. The resulting typical plot (**Figure 5d**) indeed shows reentrance and second transition at a higher concentration of ions exactly as observed in experiments.^26^ As an important control we explore a hypothetical situation where ions charges are abrogated [A=0 in **Eqs. (3)-(5)**]. As seen on **Figure 5e** the reentrant LLPS transition at low salt concentration disappears while the transition at higher salt remains unaffected exactly as observed in simulations (**Figure 1**).

Finally, to get a deeper insight into the origin of the low and high salt transitions we explore temperature dependence of both (**Figure 5f**). As can be seen clearly, in a condensed phase formed at lower concentration of salt only the first reentrant transition occurs at low temperatures but the second transition at high salt appears at higher temperature and becomes stronger as temperature increases while the low salt transition becomes weaker and finally disappears as temperature increases. At intermediate temperatures both transitions occur in their respective ranges of salt concentration. From that we conclude that first transition is driven by energetics of FUS-ion and ion-ion interactions and is opposed by unfavorable entropy of redistributing of ions inside/outside the condensate while second transition is clearly driven by entropy that favors more uniform distributions of all components of the complex solution inside the condensate at high salt concentrations.

In summary, the predictions from analytical theory are:

1. The reentrant transition at low ionic strengths is driven by ionic charges whereby FUS monomers effectively interact through the medium of highly correlated salt ions. In this regime ion concentration inside the condensate exceeds that of the bulk solution.
2. At a higher ionic strength charge correlations become short-range and FUS condensate dissolves.
3. At even higher (non-physiological) ionic strengths another FUS condensation transition, driven by entropy of mixing, occurs. In this regime ionic charges do not play a crucial role, and the solvent components redistribute in the opposite direction, making concentration of ions inside the condensate lower than in the bulk. This transition is akin to collapse transition in a mixed solvent studied in *Ref*. 56 by Xia *et al*.

## Discussion

Studies of charged polymers (polyelectrolytes and polyampholytes) in solutions containing salts have a long history.^34-36, 61-64^ Discovery of membrane-less organelles in living cells formed by IDPs undergoing LLPS^26^ reignited interest in biopolymer condensates in physiologically relevant environments.^65^ Statistical mechanical analysis using field-theoretical tools from polymer theory^66, 67^ provided detailed phase diagrams for specific IDPs enriched in charged amino acids.^37^ In these cases, salts affect the phase state of a polyelectrolyte by screening electrostatic repulsion between its charges giving rise, in some cases, to LLPS. However, coacervation experiments did not reveal salt dependent re-entrant transition.^68^ In a similar vein, coacervation of polyelectrolytes of opposite charges appeared to be strongly salt-dependent as electrostatics is a driving force in these situations.^62, 68^

Wessen *et al*. explored the phase behavior of net neutral or nearly net neutral polyampholytes by using polymer field theory and random phase approximation (RPA) methods.^59^ They found that certain polyampholyte sequences can exhibit reentrant phase transition with respect to ionic strength of the solution. Although there is net electrostatic neutrality in polyampholytes, the charge content is high, unlike the FUS type sequences of mostly neutral, uncharged amino acids which have a very low fraction of positive and negative charges (*f*_+_ *and f*_*−*_). This gives rise to fundamental difference in phase behaviors is as follows.

Polyampholytes can form condensates without the presence of any ions (that is at 0.0 M salt) as shown in **Figure SG**, consistent with Wessen *et al*.^59^ In contrast FUS-LC type uncharged sequences cannot form condensates without salt ions present. In case of polyampholytes, the presence of salt screens attractive interactions between charges leading to de-condensation transition at higher salt concentration. In case of FUS-LC type uncharged IDP salt ions *drive* condensation. In the case of polyampholytes, the driver of condensation is the charges on the IDP. Turning off the charges of the salt ions does not dissolve the polyampholyte condensate, which is also different from the behavior shown by FUS-LC condensate. *Apparently, the physics of condensation transition in globally neutral polyampholytes and salt induced transition in uncharged IDPs are drastically different*. A further discussion of this point is presented in the **Supporting Information** (Section 9.

Recent experiments showed that largely uncharged IDPs such as FUS undergo a set of complex reentrant transitions in a broad range of salt concentration.^33^ The authors of Ref. 33 first carried out a series of atomistic simulations to demonstrate that different forces are dominant at different salt concentrations. While insightful these studies do not provide a clear mechanistic explanation of the multiple transitions that FUS and other uncharged IDPs undergo in a broad range of salt concentrations with multiple instances of reentrance. We further note that Krainer *et al*. employed the implicit solvent CG force field (no bead representation for ions and water) to simulate condensates, however by *tweaking the relative contribution of electrostatic and hydrophobic interactions among amino acid pairs*, according to the salt concentration, based on their atomistic PMF results.^33^ Therefore, in its original form with Debye-Huckel electrostatics and no explicit ions/water, the CG model they used would be incapable of capturing the reentrance. On the other hand, MARTINI can capture the reentrant without the need for salt-concentration specific parameterization, probably due to its explicit description of ions and water. The latter also reveals the spatial distribution of ions which was not captured through implicit ion/solvent simulations.

In this work we used a combination of theoretical and computational approaches to reach mechanistic understanding of the complex reentrant phase behavior of uncharged IDPs such as FUS at different salt concentrations. The mechanistic picture emerging from these analyses is as follows. At low salt concentrations LJ interaction between an uncharged FUS monomer and a salt ion creates local density perturbation creating a density gradient in a highly correlated ion plasma medium that extends up to Debye length. Given that interactions between ions and FUS monomers are energetically favorable due to LJ attraction the excess of ion density created by one FUS monomer which propagates up to correlation length in the ion media serves as an effective energetic funnel creating a long-range attractive potential for another FUS monomer(s). This effective long-range attraction between uncharged FUS monomers is illustrated in **Figure 2a** which shows the PMF between two FUS monomers in the ionic media derived from MARTINI simulation. In the full model where ion charges are intact the PMF shows effective long-range attractive interactions that extend up to 12 nm length scale at physiological salt concentration (blue line of **Figure 2a**). When ion charges are abrogated or only one kind of ion is present this effective attractive potential disappears (red and green lines on **Figure 2a**).

This mechanism is conceptually similar to polymer-salt-induced condensation, often referred to as ψ-condensation.^57^ *ψ*-condensation refers to collapse of DNA in the presence of a uncharged polymer (PEO) and monovalent salt in solution.^69^ In *ψ* -condensation, DNA-induced perturbation of density of PEO propagates at longer scales due to correlated fluctuations in polymer solution creating an effective long range potential well for DNA^57^ – a mechanism similar to the one described here where the role of correlated medium is played by ions.

As salt concentration increases beyond physiological levels, screening effects become increasingly significant, leading to a suppression of long-range attractive interactions and, ultimately, dissolution of the condensate. Our results show that this dissolution occurs around between 1M–2M salt concentration. However, an intriguing phenomenon emerges at even higher ionic strengths, where LLPS reappears beyond 2M salt concentration, marking the second reentrant transition. Unlike the condensation mechanism at low salt, phase separation in this regime is not driven by FUS-salt interactions but rather by an ion-exclusion mechanism akin to depletion forces. The FUS condensate expels ions from its interior, allowing the system to regain phase separation despite the high salt environment. The second LLPS transition at high salt concentration is an entropy driven phenomenon which disappears at low temperature as predicted by the theory (see **Figure 5e**). CG simulations show the same behavior: while transition at low salt is crucially dependent on ion charges the second transition at high salt is insensitive to the abrogation of charges of salt ions suggesting that it is driven mainly by LJ energy of association between FUS and salt and, crucially entropy of all components in the condensate.^56^

This is a striking departure from the behavior observed at low ionic strength, underscoring the nontrivial role of electrostatic correlations and entropic forces in governing phase behavior in IDPs with a low fraction of charged residues. A key insight from our study is that the mechanism of LLPS at low and high salt concentrations is fundamentally distinct, even though both regimes exhibit phase separation. This distinction is evident in the differences in ion distribution, pair interaction energies, and overall condensate morphology. The ψ-condensation-like mechanism at physiological salt concentrations leads to a system where ions are deeply embedded in the condensate, while at extreme salt conditions, the driving force shifts towards an entropic phase separation driven by ion exclusion. These findings not only provide a microscopic understanding of the reentrant LLPS of FUS but also suggest a generalizable framework for other IDPs with low charge density.

Overall, our study provides a comprehensive statistical mechanical view of mechanisms of LLPS in water-salt solutions where theoretical predictions are tested and verified by molecular simulations. By unraveling the mechanistic details of LLPS across different ionic environments, we not only explain recent experimental observations but also establish a broader paradigm for understanding phase separation in biologically relevant uncharged systems.

## Acknowledgements

This work is supported by the National Institute of Health (Grant R35 GM139571). SM and ES thank the ‘Faculty of Arts and Science Research Computing’ (FASRC) at Harvard University for computational resources.

## Supporting information *for*

### 1. Detailed steps of the analytical theory

Consider the phase separated solution of FUS. It occupies volume V lattice cites (together with ions) while the rest of the solution in the dilute phase occupies the remaining volume V_0_-V. The first region we call “inside” (in) and the polymer free region we call outside (“out”). We do not make distinction between Na^+^ and Cl^−^ ions (same interactions with polymers for simplicity). We will use Flory-Huggins approximation to present free energy of the system.

We consider a general two component phase separated system with volume of dense phase V, total volume V_0_, total number of FUS chains, ions, and water molecules and the same in the dense phase correspondingly (we mark molecules in dense phase with symbol “in”, molecules in dilute phase symbol “out” and total with symbol “0”) 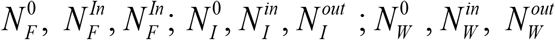. Free energy of the whole system is then:

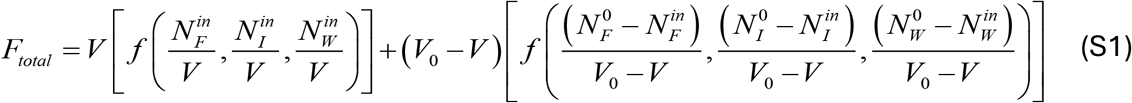

Adding condition of incompressibility and assuming that monomers of each type are of the same volume *v*_0_

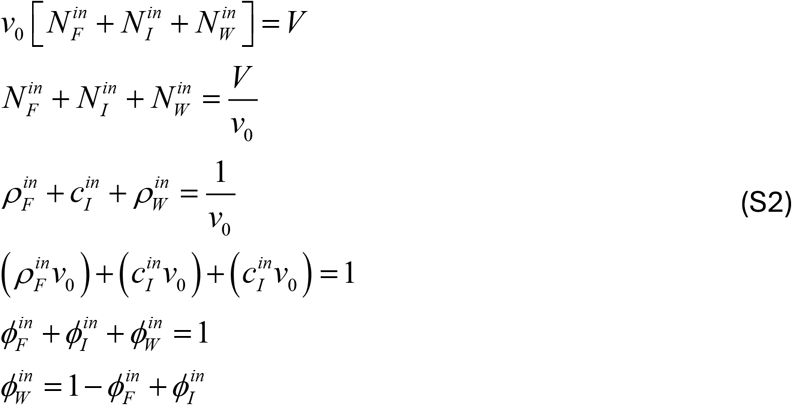

where *L* is the polymerization index of FUS. We switched from densities to volume fractions using the relation *ϕ* = *ρν*_0_ or *ϕ* = *cν*_0_. Equivalently we can derive the same equation for outside of the dense phase

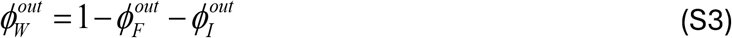

Equilibrium between dense and diluted phases of FUS calls for equal chemical potentials of FUS molecules between phases and equal chemical potentials of salt ions between the dense and diluted phases. In addition, the condition of equal osmotic pressure determines the partitioning between the phases, i.e. determines the volume of the dense phase V:

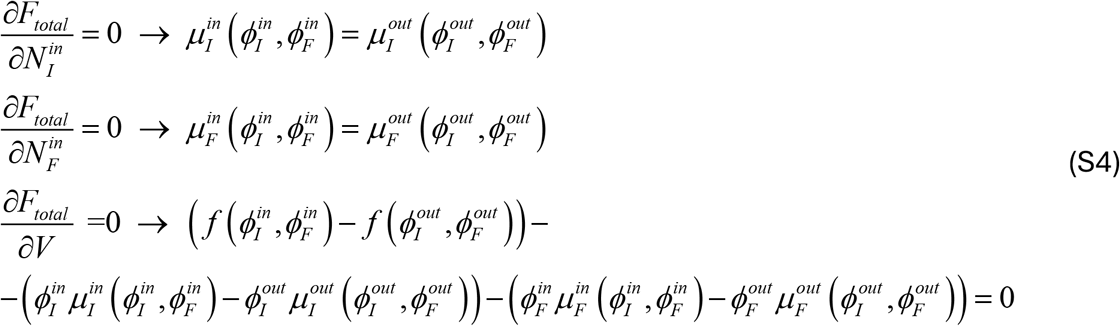

The condition of conservation of total amounts of FUS and ions are:

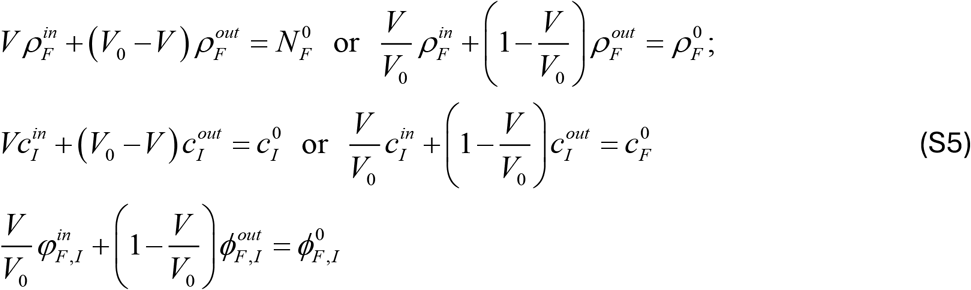

Where 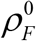 and 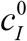 are initial total concentration of FUS and ions respectively.

Excluding water using the incompressibility conditions leaves us with effective free energy in the Voorn-Overbeek approximation^1^ (**Eq. (2)** of the Main text) which treats polymers in the FH approximation and ion free energy in the Debye-Huckel approximation. As pointed out in the main text 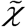 in Eq.2 are effective interactions between FUS and Ions – effective because water interactions are excluded through incompressibility conditions.

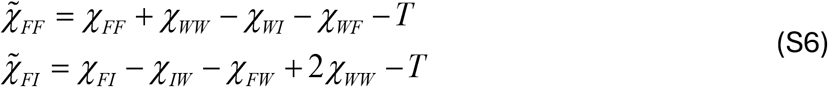

*F*_*DH*_ is free energy of ion-ion interactions in the Debye-Huckel approximation which will be detailed later.

In the regime where both FUS and ion concentrations are low 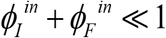 we can omit water entropy term in **Eq. (2)** of Main Text. Denoting 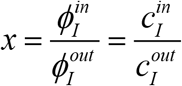 and substituting into **Eq. (S6)** we get for the chemical equilibrium for the ions (see **Eq.3** of main text and accompanying discussion)

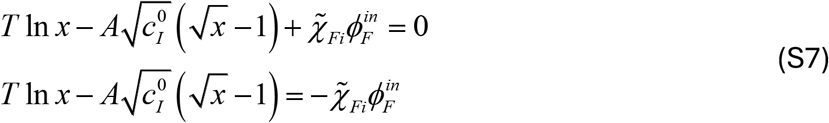

Maxima of the curves shown in **Figure S1(a)** are reached at

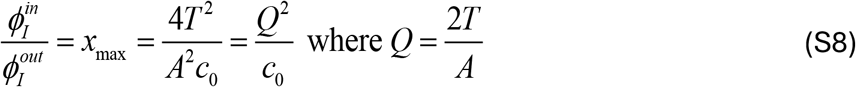

Maximal value of 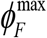 possible is given by:

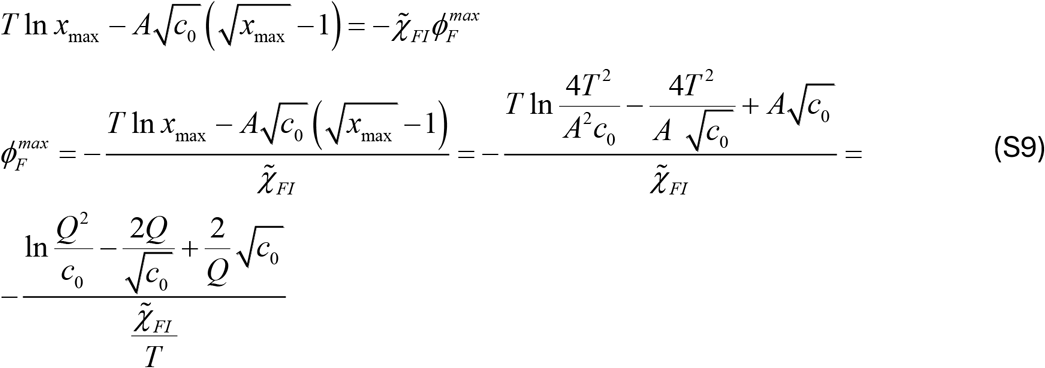

The plot of 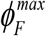 vs salt concentration is shown in **Figure S1(b)**

**Figure S1.**
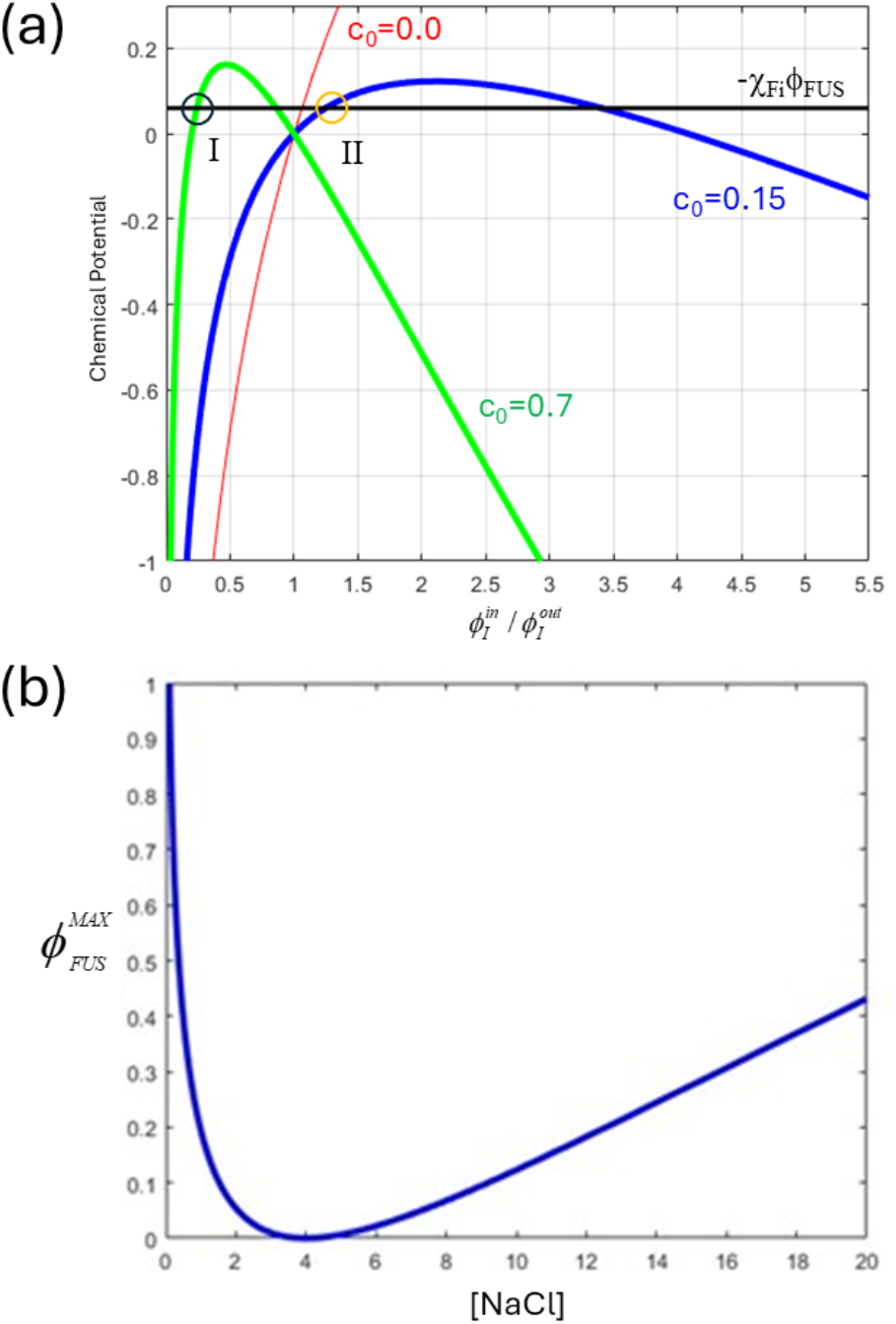
(a) Graphical solutions of Eq. (S7): The red line: no salt charges, (just inert crowders instead of charged ions), the blue line 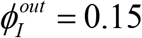,the green line 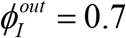, the black line constant RHS of −𝒳_*Fi*_ϕ_*FUS*_. The intersections marked with ‘I’ and ‘II’ are the stable solutions. **(b) Maximal density (volume fraction) of FUS at which Eq.(SG) has nontrivial solutions** as a function of [NaCl] concentration (in arbitrary units). At higher volume fractions of FUS equilibrium is possible only at 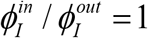 and ϕ_*FUS*_ = 0 which is the case at intermediate salt concentrations where non-trivial equilibrium is achieved at very low FUS volume fractions indistinguishable from random coil leading to de-collapse upon increase of [NaCl] and collapse again at much higher [NaCl].

It is clear from this plot that at low salt concentration the chain can collapse (provided that free energy of the collapsed state is lower than random coil, see below), at intermediate concentrations it can be only random coli and at higher concentrations the chain can collapse again as MAX density of FUS again rises. That explains, in principle two reentrant transitions. At *c*_0_ < *Q* maximum is at 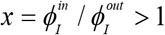.

The plots on Figure S1(a) show that for the purpose of solving the equation in the relevant range of x we can approximate the chem potential function as a parabola:

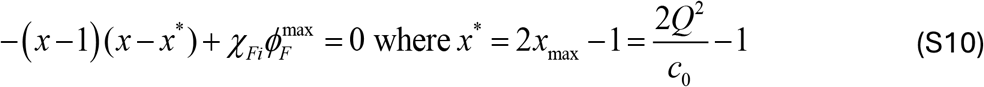

Solving for x we get:

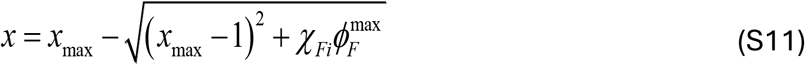

Giving us the result for concentration of ions inside the FUS polymer as

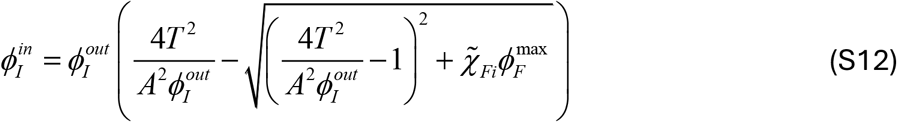

Now consider the onset of collapse i.e. 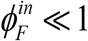. In this case we expand the term with the square-root, in Eq.(S15) and get finally:

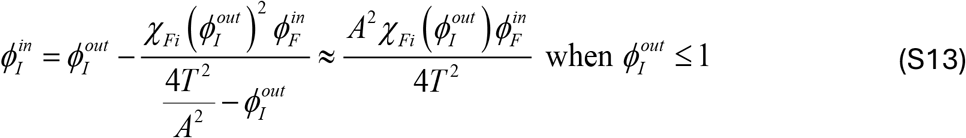

Substituting this expression for *ϕ*_*out*_ into the original FH **Eq. (2)** of main text we get

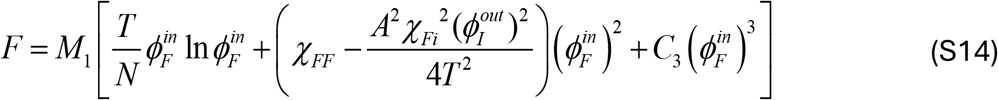

Here we see that expulsion of ions effectively leads to additional attraction between FUS monomers by renormalizing its second virial coefficient of interactions

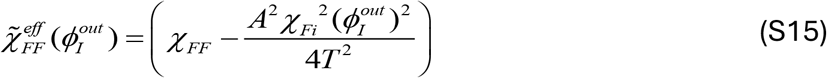

When

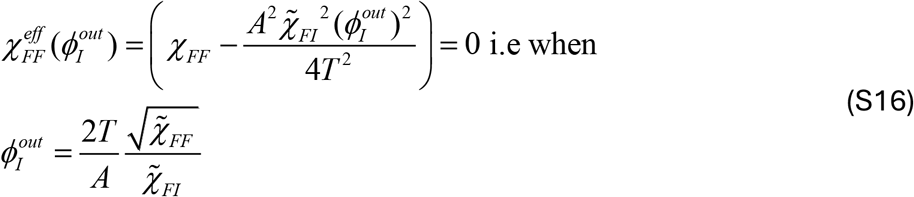

The chain collapses, i.e. it reaches an effective *θ*-point.

### 2. Sequence Analysis

**Table S1(a).**
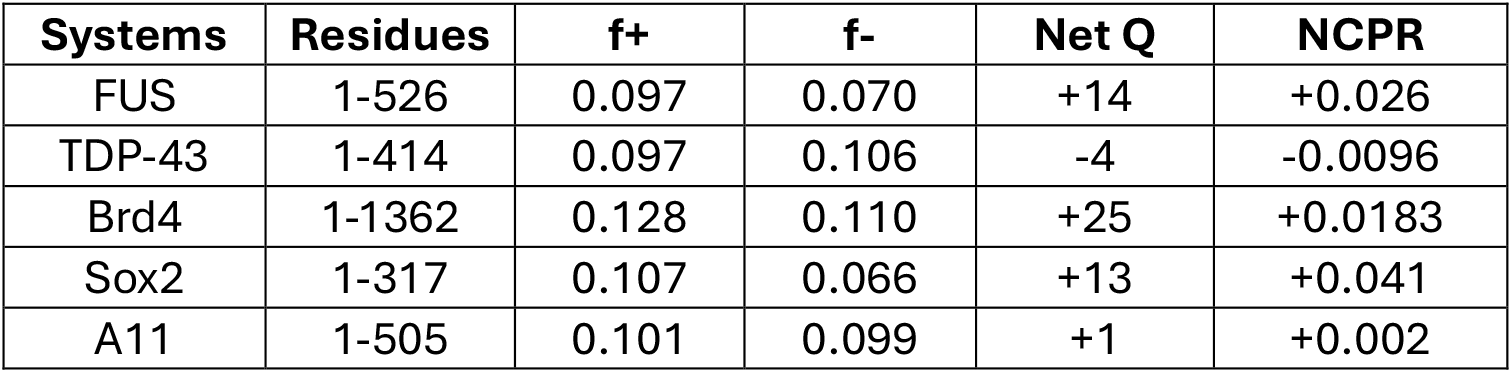
Fraction of positively charged residues (f+), fraction of negatively charged residues (f-), total charge (Net Q), and net charge per residue (NCPR) for the five full length constructs used in the experimental study by Krainer *et al*.^2^.

**Table S1(b).**
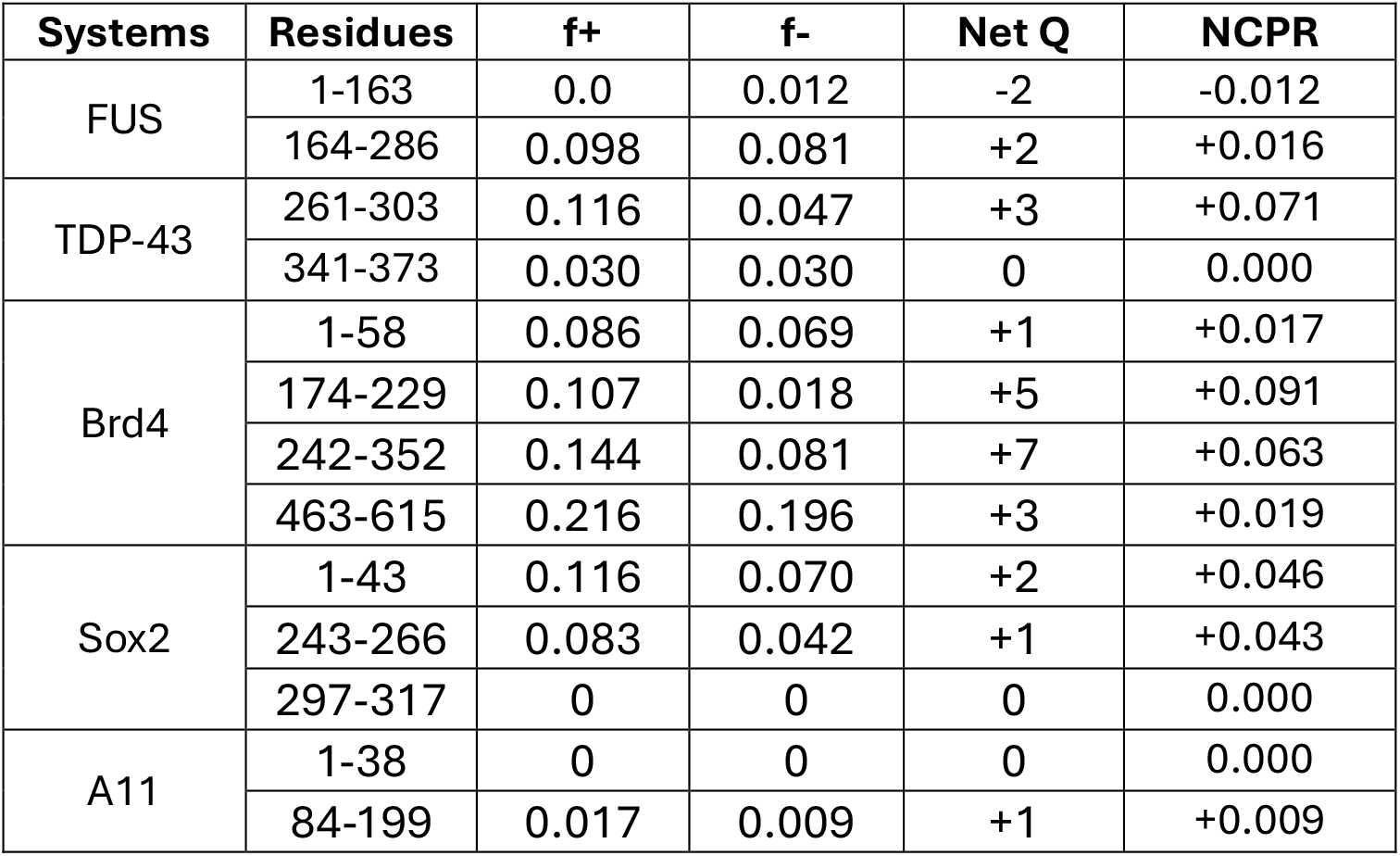
The same parameters calculated for the disordered (low complexity) droplet promoting regions of the five systems listed in Table S1(a).

### 3. Pairwise Interaction Energetics Figures

In this section, we plot the pairwise interaction energetics of different pairs namely, inter-protein (*E*_*FUS*_), protein-water (*E*_*FUS*−*W*_), protein-ion 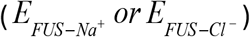, and inter-ion interactions (*EION* −*ION*); for both the original system (**Figure S2**) and a control system where the ionic charges are removed (**Figure S3**). For the latter, there is no distinction between sodium and chloride ions. Hence, instead of 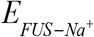 *or* 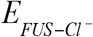 we use *E*_*FUS* −*ION*_. All the energy values are a combination of electrostatic (Coulomb) and van der Waals (Lennard-Jones) interactions, and are normalized with respect to the number of FUS chains, *N*.

**Figure S2.**
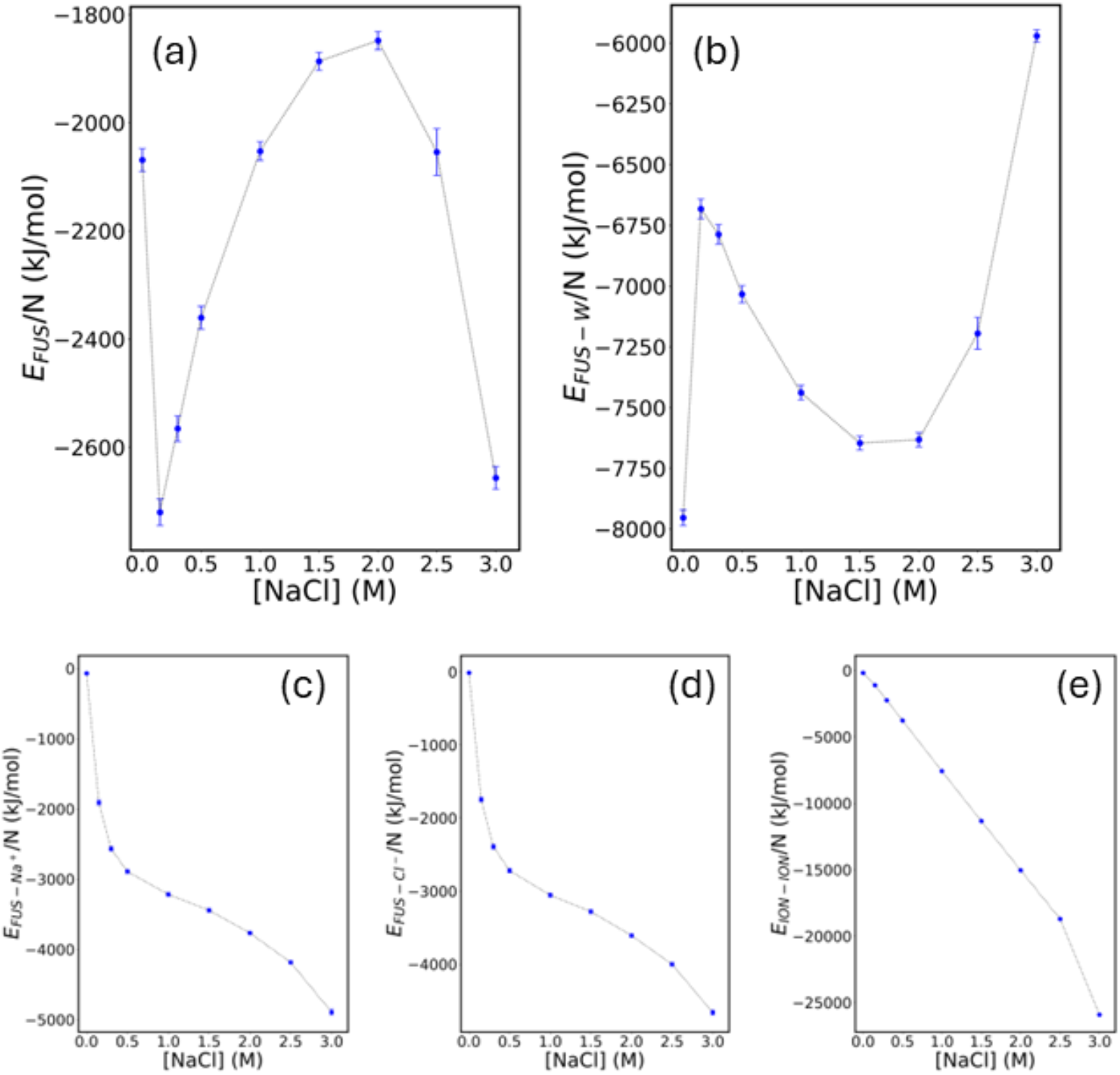
Pairwise interaction energetics against the solution’s ionic strength (when the ions carry their natural charges): (a) Inter-protein interaction sharply drops at 0.15 M [NaCl] followed by a gradual increase, reaching a maximum, and again dropping sharply at 3 M [NaCl]. The sharp decreases indicate condensation. (b) The protein-water interaction energy shows a sharp increase at 0.15 M [NaCl] followed by a gradual decrease, reaching a minimum, and again increasing at 3 M [NaCl]. (c) Protein-sodium ion and (d) protein-chloride ion interaction energetics exhibit monotonic decrease with the increasing salt concentration. (e) Inter-ion interactions also show a monotonic decrease with respect to solution’s ionic strength.

**Figure S3.**
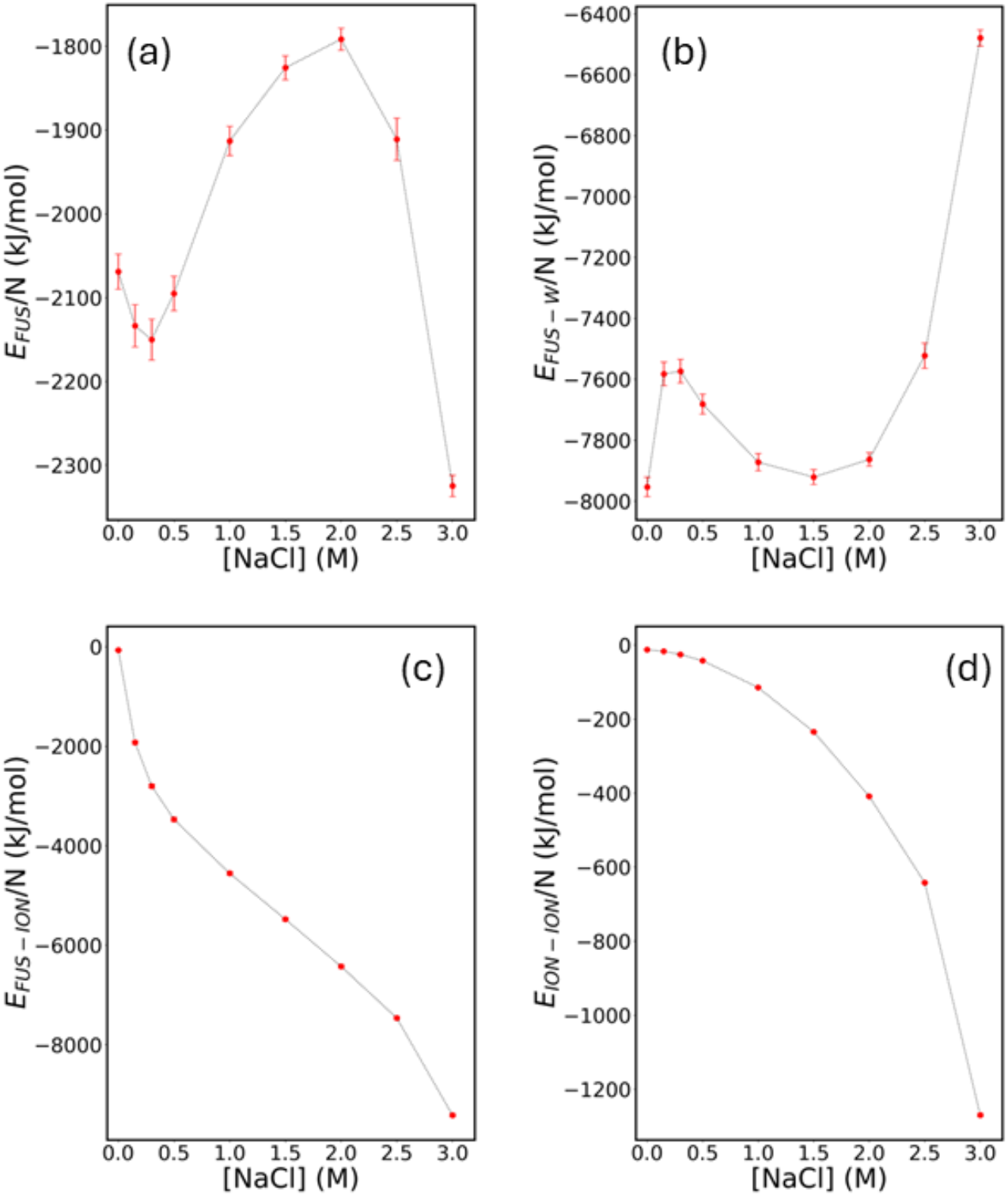
Pairwise interaction energetics against the solution’s ionic strength (when electrostatic interactions are ‘turned off’ that is, ionic charges are set to ‘0’): (a) Inter-protein interaction shows a little dip around 0.15 M [NaCl] before increasing, reaching a maximum, and decreasing sharply at 3 M [NaCl]. The initial dip indicates enthalpic stabilization to some extent but not condensation. (b) Protein-water interaction initially shows a slight increase followed by gradual decrease, reaching a minimum, and a sharp increase at 3 M [NaCl]. (c) Protein-ion interaction and (d) inter-ionic interaction show monotonic decrease with increasing [NaCl].

### 4. Condensation in System-2 Starting from a Dispersed State

System-2, where the FUS charges are removed but salt ion charges are kept intact, exhibit LLPS. In the main text (**Figure 3b**) we showed the final state of the system starting with a preformed droplet state. To rule out a possible kinetic trap we carried out additional simulation starting with a fully dispersed state and observed spontaneous LLPS within a few microseconds of coarse-grained timescale (**Figure S4**).

**Figure S4.**
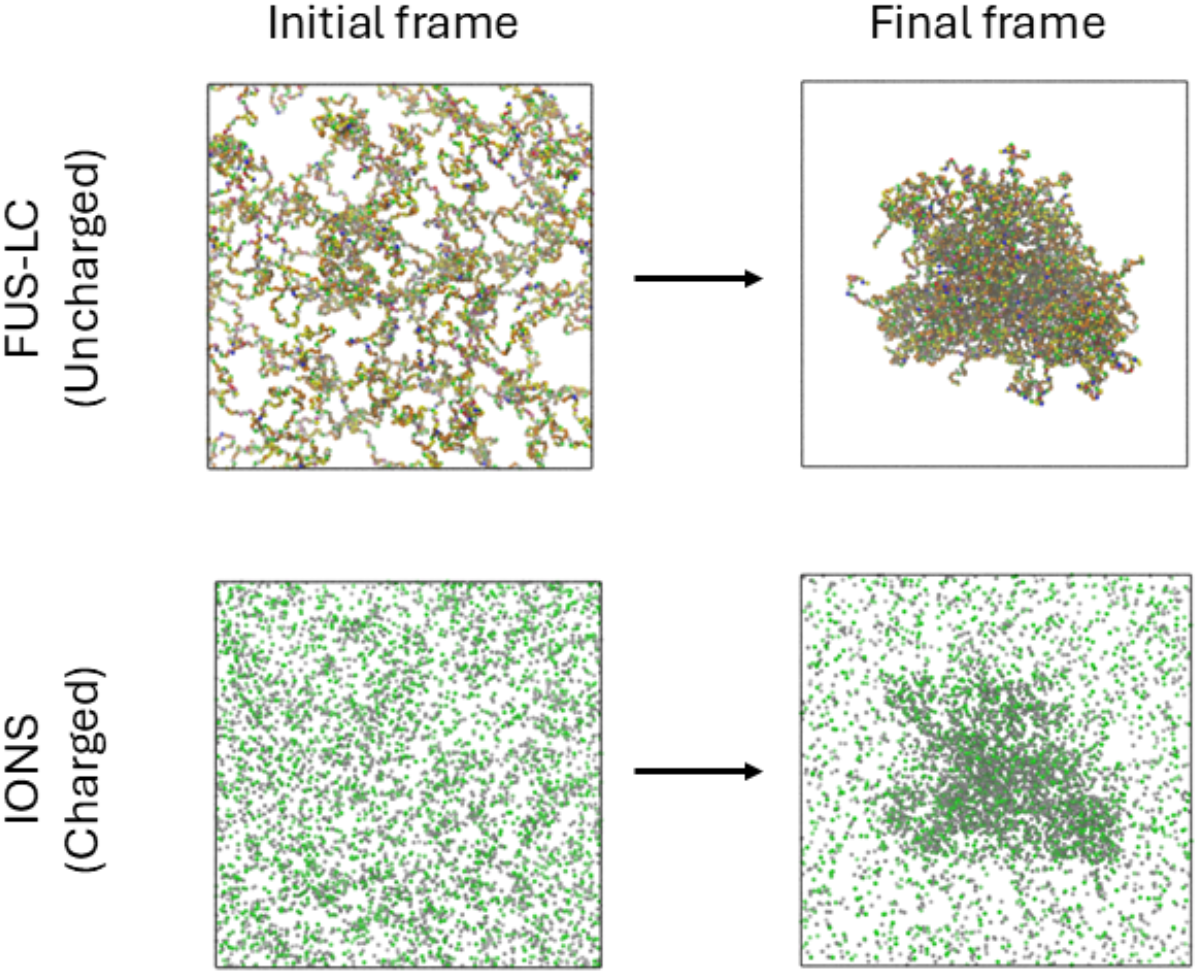
LLPS of uncharged FUS-LC with charged 0.15 M NaCl, starting from a fully dispersed state: Top panel shows the distribution of FUS chains at their initial (dispersed) and final (condensed) state. The bottom panel shows the distribution of Na+ (grey) and Cl-(green) ions for the same timeframes as above.

### 5. Comparison between MARTINI-3 and Madrid-201G model contact ion pairs

We have simulated NaCl ions in water at two different high salt concentrations, namely 1.50 M, and 3.00 M. At T=298K and p=1 bar, from classical MD simulations, we have obtained contact ion pair (*nCIP*) from the partial radial distribution function, g(r) between Na + and Cl − as follows:

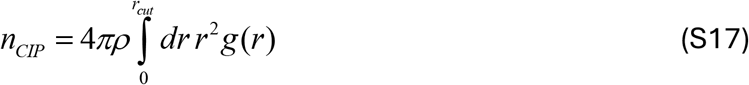

where *r*_*cut*_ is the cut-off distance up to which we integrate g(r) and *ρ* is the number density of either Na^+^ or Cl^−^. We compare the MARTINI-3 results with one of the best available forcefields for ionic solution (Madrid-2019 with TIP4P/2005 water model)^3^ that predicts NaCl solubility close to 6 M. For comparison purposes, we set *r*_*cut*_=0.6 nm which is close to a minimum in g(r). The results are tabulated below:

**Table S2.**
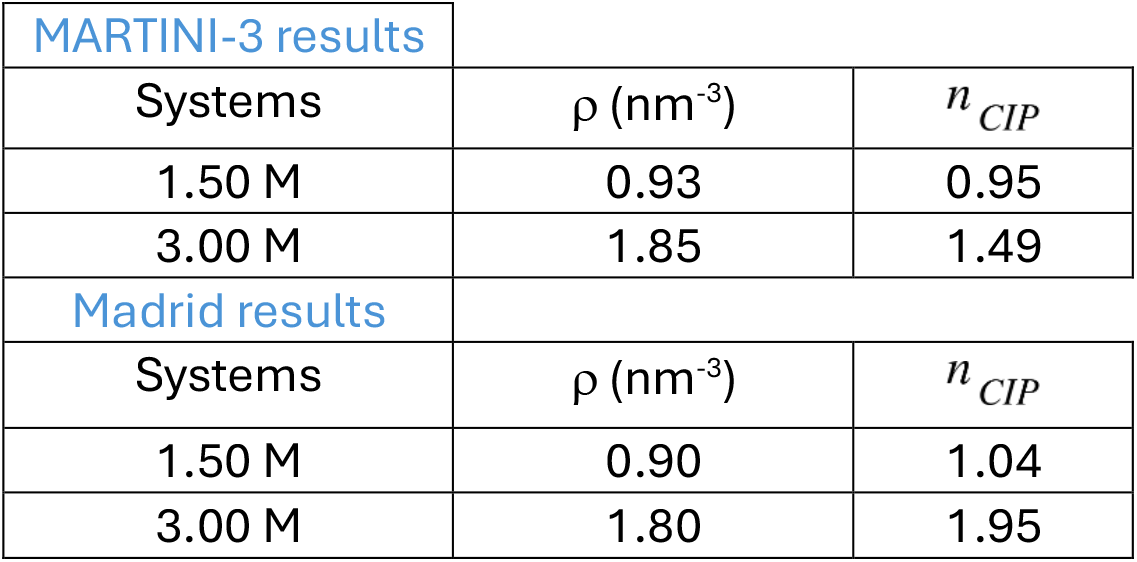
Contact ions pairs obtained from MARTINI-3 (coarse-grained) and Madrid (atomistic) force fields for two different high ionic strengths.

From **Table S2**, we notice that the number densities (after NpT simulation) obtained from MARTINI force-field, on average, are only 3% higher than those obtained from the atomistic force field. For the MARTINI model, the values of *n* _*CIP*_ are even lower than those of Madrid model. A higher value of *n* _*CIP*_ denotes an increased ‘crowding’ of cations (or anions) around anions (or cations). We provide the radial distribution plots below (**Figure S5**), where the solid lines denote Atomistic force field and dashed lines denote coarse-grained force field.

**Figure S5.**
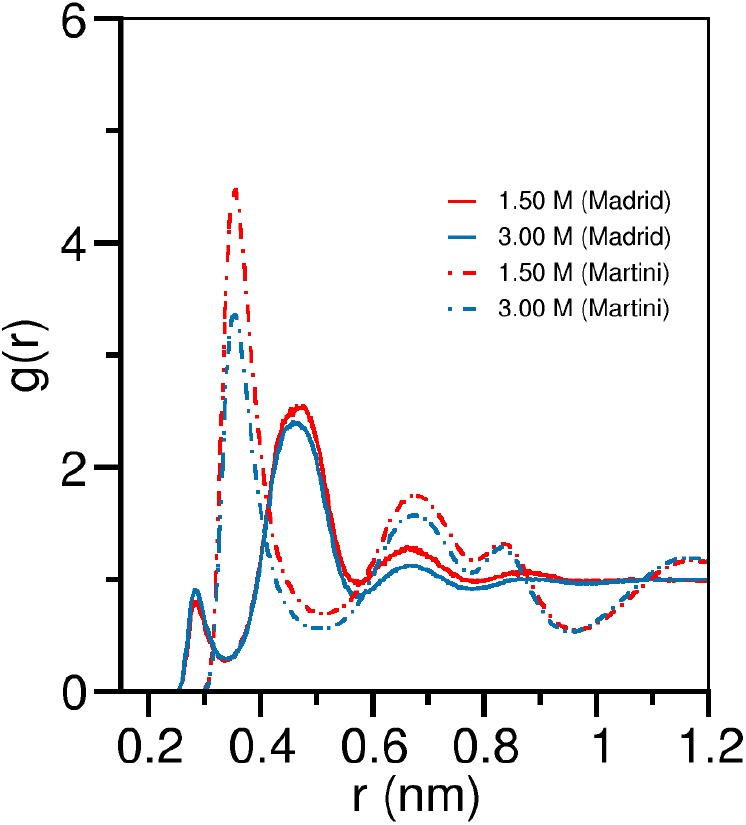
Radial distribution function (RDF) comparison between atomistic and MARTINI models: RDF of Cl-ions with respect to Na+ ions in two different salt concentrations, by using Martini (coarse-grained) and Madrid (atomistic) models.

The two peaked fine structures of g(r) in the atomistic force field (centered around 0.28 nm and 0.47 nm) are merged to one peak structure (centered around 0.36 nm). This can be attributed to the difference in the van der Waals radii (σ) of ion and water beads. For example, in the atomistic force field 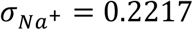 and 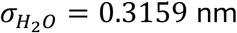; whereas in the MARTINI coarse-grained description 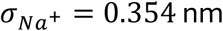 and 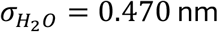. As a result, the finer peak structures in g(r) disappear and a ‘coarser’ peak appears in between. Nevertheless, the value of *n*_*CIP*_ remains comparable between the two force fields.

### 6. LLPS of full-length FUS

In the main text, we discussed the reentrant LLPS of the low complexity domain of FUS. Here, we provide the LLPS propensity of full-length FUS at three different salt concentrations, namely, 0.15 M, 1.5 M, and 3.0 M. We find that, the full-length FUS exhibits condensation at 0.15 M and 3.00 M, but not at 1.5 M. This corroborates well with the observation with the LLPS of FUS-LC only. However, the phase boundaries may change/shift if one compares the phase behavior of the full-length FUS and the LC region. The comparison can only be in a qualitative sense.

**Figure S6.**
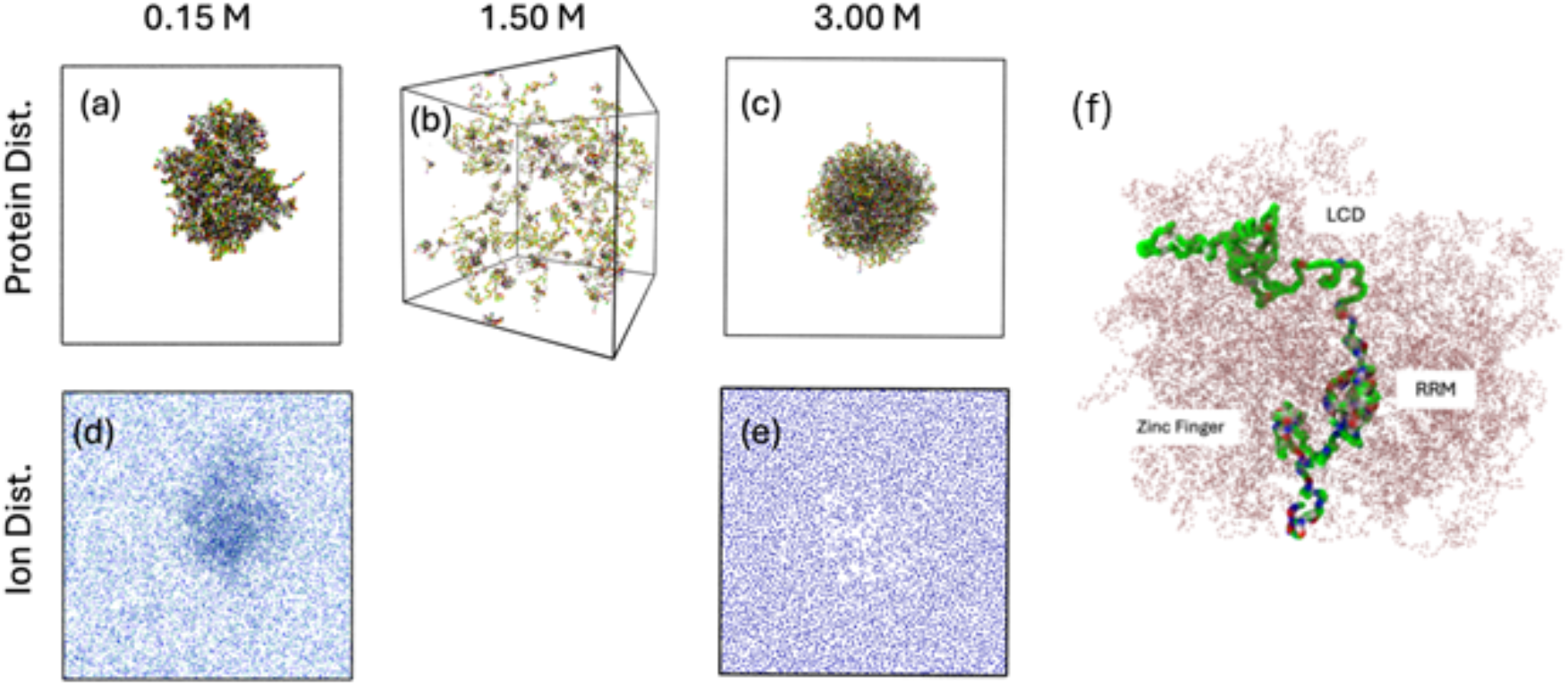
MD simulation of Full-length FUS: (a), (b), and (c) are representative snapshots of the system after 3 ⍰s MD simulation for three different salt concentrations: 0.15 M, 1.5 M, and 3.0 M respectively. (d) and (e) are the spatial distribution of ions inside the box at the exact same time-point as the condensate snapshots. The observations by simulation of the FUS-LC domain are valid when full length FUS is used. (f) Magnified view of the condensate in panel ‘a’ where the conformation of a single full-length FUS is highlighted.

### 7. LLPS of FUS-LCD at less than 0.15 M NaCl

To comment on the threshold salt concentration of the LLPS of FUS-LCD, we have carried out additional coarse-grained simulations at [*C*_*ion*_] = 0.05 M and 0.10 M for 5 μs starting form a completely dispersed state. We observe LLPS at [*C*_*ion*_] = 0.10 M but not at [*C*_*ion*_] = 0.05 M (**Figure S7**). This suggests that the threshold lies in between 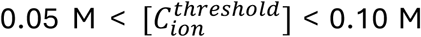. We reiterate that these results are with FUS-LCD and the threshold might differ for full length FUS. In **Figure S7(a)** we show the time evolution of the radius of gyration of the entire protein system and in **Figure S7(b)** we show the distribution of number of clusters.

**Figure S7.**
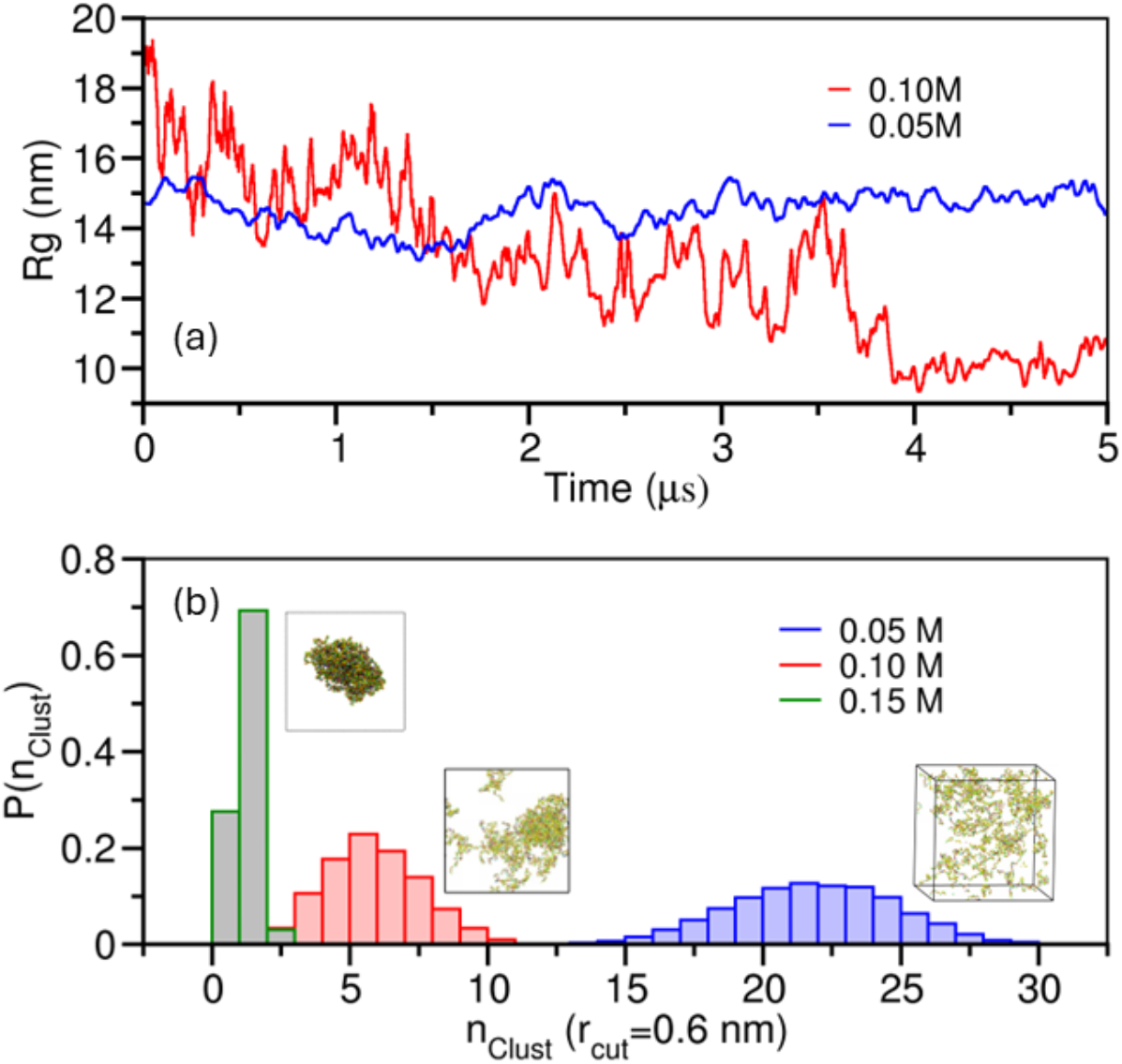
LLPS propensity of FUS-LCD at salt concentrations lower than 0.15 M: (a) Time evolution of the radius of gyration (Rg) of all the proteins. At 0.05 M the Rg shows no sign of decrease along the trajectory whereas at 0.10 M the Rg significantly drops around 4 ms. (b) The distribution of numbers of clusters in the system shows right shifted distribution at 0.05 M indicating the presence of several smaller clusters. At 0.10 M the distribution shifts to the left indicating the onset of LLPS (snapshot beside the distribution) and as a reference, the sharply peaked distribution is at 0.15 M.

### 8. Validation from atomistic simulations

we have performed atomistic simulations with CHARMM36 forcefield and TIP3P water model. We have utilized the back-mapping algorithm to convert the equilibrated coarse-grained condensates into atomistic resolution (for details please see https://cgmartini.nl/docs/tutorials/Martini3/Backward/). We have additionally modified the Lennard-Jones parameters of Na^+^ and Cl^-^ions following the work of Yagasaki *et al*.^*4*^ who obtained a solubility of 6.1 ± 0.30 mol/kg for NaCl in TIP3P water model.

**Figure S8.**
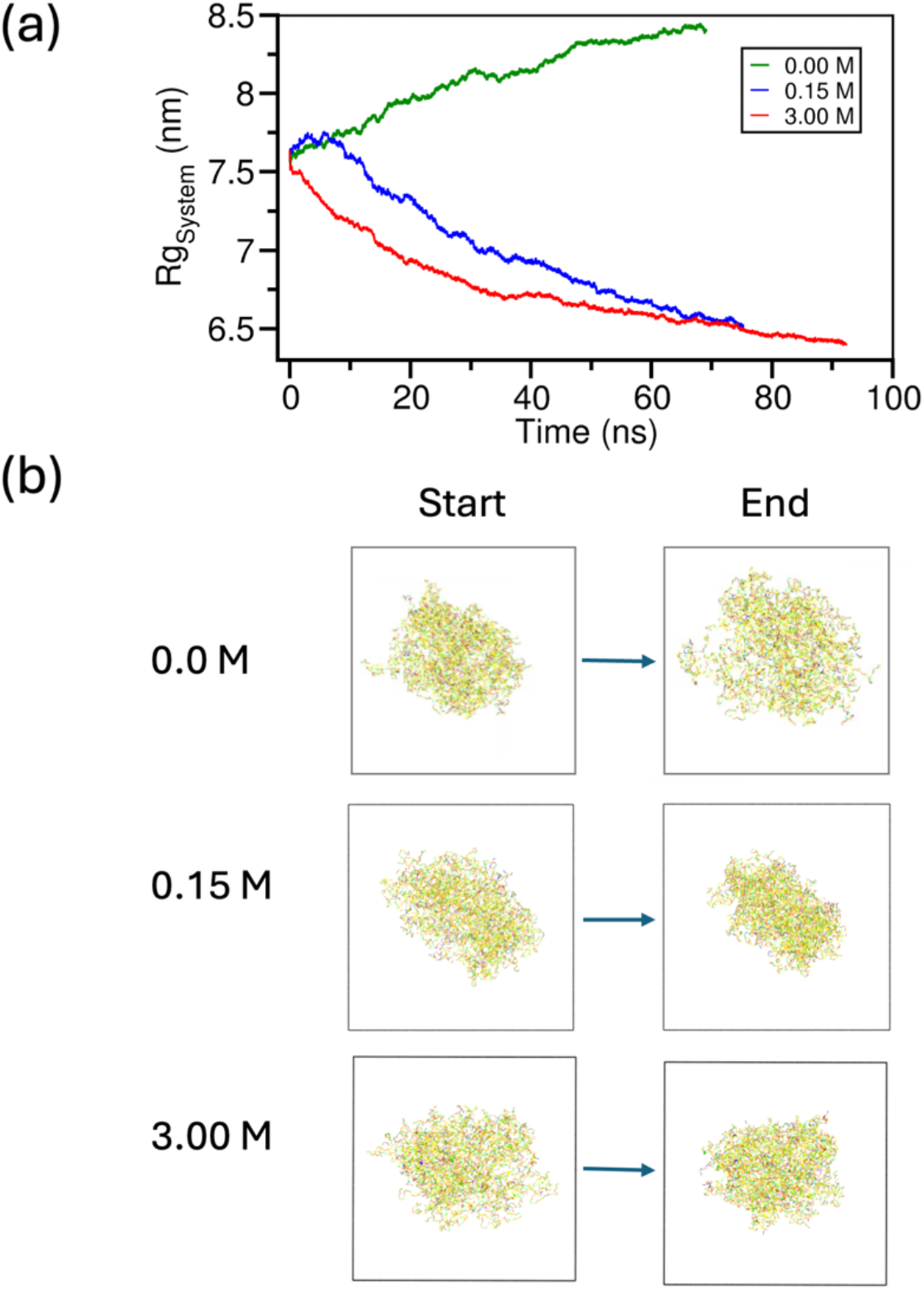
Atomistic simulation of a preformed droplet at three salt concentrations. (a) Time evolution of Rg of the protein system during atomistic MD simulation shows a gradual increase in Rg for 0.0 M salt and a gradual decrease in Rg indicating compaction of the condensate for 0.15 M and 3.0 M, compared to its backmapped configuration from MARTINI. (b) Snapshots of the systems before and after atomistic MD simulations at three different salt concentrations showing swelling for 0.0 M and compaction for 0.15 M and 3.0 M during the atomistic simulation.

We started the atomistic simulations with a preformed droplet state and run the MD simulations up to 100 ns at 0.0 M, 0.15 M, and 3.0 M to check its stability. We note that starting from a dispersed phase would have been computationally challenging to simulate the timescale and length scale of interest. From the atomistic simulations, we observe that at 0.0 M salt, the Rg of the system shows a gradual increase and the condensate swells from its initial configuration. On the other hand, at 0.15 M and 3.0 M Rg of the system shows a gradual decrease indicating compaction of the condensate. However, at this short time scale (∼100 ns) we can neither obtain a fully dispersed state starting from a condensed phase, nor obtain a condensate from a dispersed phase. Nevertheless, these results are indicative of the fact that the condensate is stable at its atomistic description at 0.15 M and 3.0 M.

### 9. Differences with polyampholyte phase separation

Wessen *et al*.^*5*^ explored the phase behavior of neutral or nearly neutral polyampholytes by using polymer field theory and random phase approximation (RPA) methods. They found that certain polyampholyte sequences can exhibit reentrant phase transition with respect to ionic strength of the solution (**Fig. 4a** and **Fig. 5** of Wessen *et al*.). They performed field theoretical simulations to show that (**Fig. 6a** of Wessen *et al*.) a polyampholyte with γ < 2 can exhibit salt dependent re-entrant whereas others with γ ≥ 2 cannot exhibit reentrant transition, in the chosen parameter space (1 being the relative change in volume upon condensation). However, there are several key differences in the physics of condensation between theirs and our systems of interest, as listed below:

1. We note the polyampholyte sequences that are *net neutral with a high fraction of positive and negative charges (f*_+_ *and f*_−_) are different from *IDR sequences that are also (nearly) net neutral but are composed of mostly uncharged residues with low values of f*_+_ *and f*_−_ (such as FUS-LC, TDP-43, etc.). Below is a comparison:
2. Because of point #1, the physics of condensation changes completely. To demonstrate this, we have additionally simulated a polyampholyte sequence (**sv10**) with MARTINI-3 force field parameterized for polyampholytes.^6^ We chose ‘sv10’ shown in Figure.S9 as this sequence was shown to exhibit re-entrance in the paper by Wessen *et al*. We find that the polyampholyte can form condensates without the presence of any ions (that is 0.0 M salt) as shown in **Figure SG**; whereas FUS-LC type sequences cannot (**Figure 3** in main text). For the latter, the presence of additional salt is required to drive the condensation. In the case of polyampholytes, the driver of condensation is the charges on the IDR. Turning off the charges of the ions does not dissolve the polyampholyte condensate, which is also different from the behavior shown by FUS-LC condensate.
3. Since polyampholytes can form condensates at 0 M salt, the re-entrant phase diagrams are different than FUS-LC type ‘nearly uncharged’ sequences. A simple schematic comparison is drawn in **Figure S10**. Panel (a) shows a representative salt concentration dependent phase diagram for FUS-LC type ‘nearly uncharged’ IDP sequences where there are 4 distinct regions, out of which α, γ are ‘No LLPS’ zones; and *β*, δ are the ‘LLPS’ zone. Along the [salt] axis, the system can exhibit three transitions as shown by purple arrows, making it a double-reentrant phase transition.

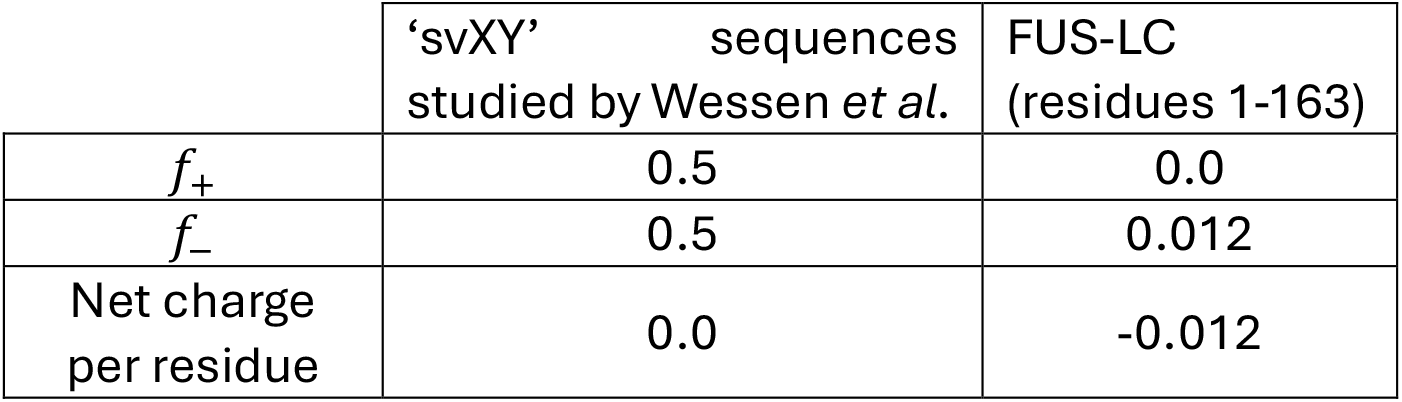

|  | 'svXY' sequences<br>studied by Wessen *et al.* | FUS-LC<br>(residues 1-163) |
| --- | --- | --- |
| $f_+$ | 0.5 | 0.0 |
| $f_-$ | 0.5 | 0.012 |
| Net charge<br>per residue | 0.0 | -0.012 |

**Figure S9.**
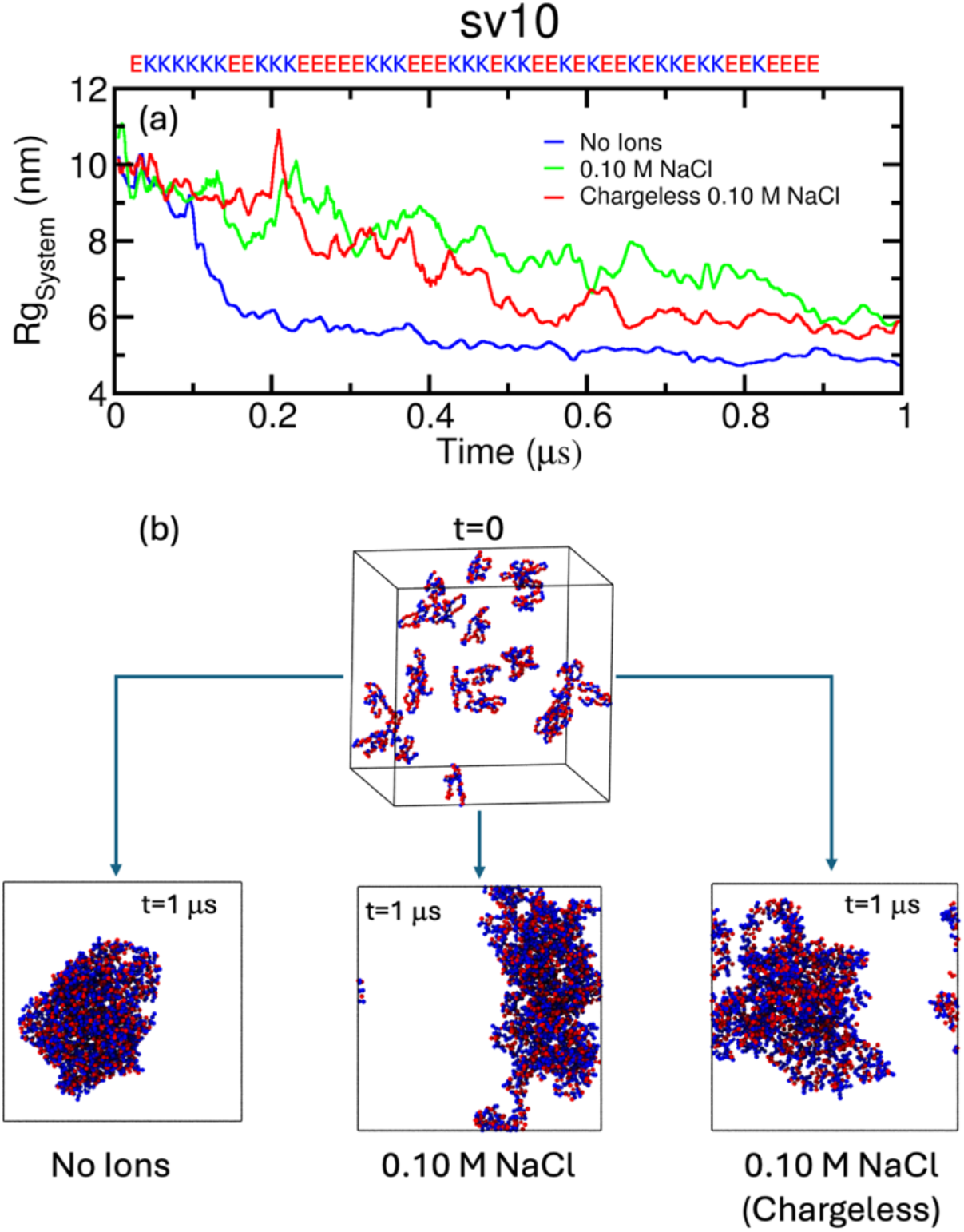
Phase separation of ‘sv10’ polyampholyte: (a) Time evolution of Rg of the polyampholyte system from 0 to 1 μs. For the system with no ions, Rg shows a sharp decrease around 150 ns indicating fast self-condensation without the help of external ions. For the 0.10 M charged/uncharged NaCl system the decrease in Rg is more gradual. (b) Representative snapshots of the sv10 polyampholyte system at t=0 and t=1 μs for three systems. The condensate is more compact when no ions were present, compared to 0.10 M NaCl charged/uncharged systems.

On the other hand, panel (b) shows a typical re-entrant phase diagram for polyampholyte sequences, that are charged but with a net neutrality. Here one finds three distinct regions: *β*′, δ′ are the ‘LLPS’ zones and γ′ is the ‘No LLPS’ zone. The system can make two transitions along the [salt] axis, making it a single-reentrant phase diagram.

**Figure S10.**
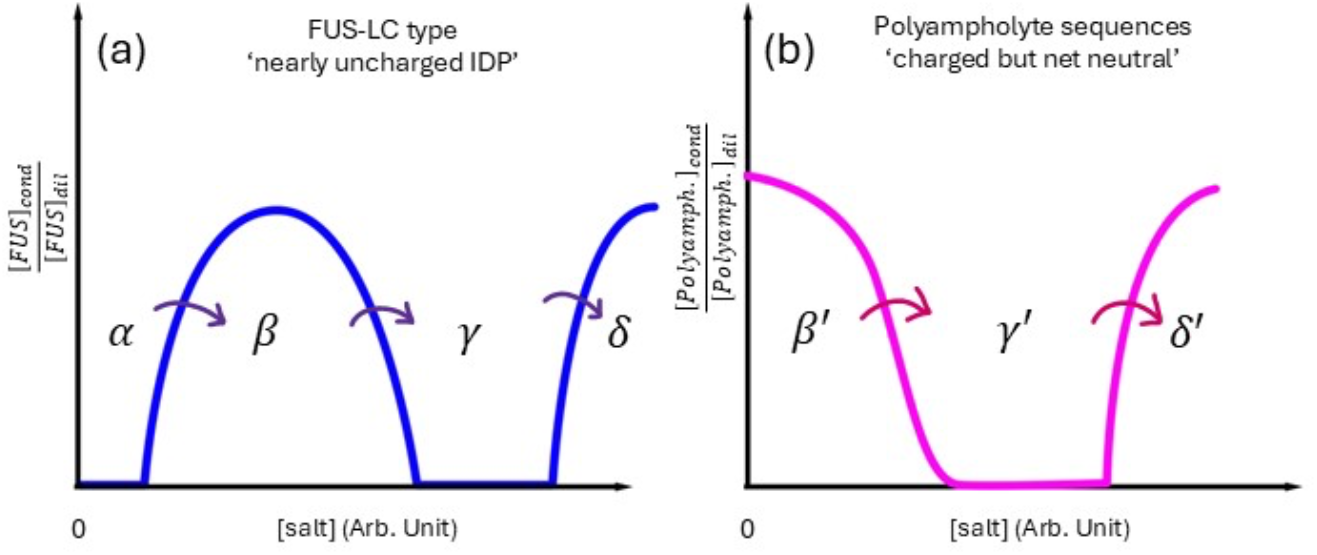
Schematic phase diagrams of (a) FUS-type nearly uncharged IDPs and (b) polyampholyte systems, against ionic strength.

