## Supporting Information for "The Origin of the Ionic-strength Dependent Reentrant Behavior in Liquid-Liquid Phase Separation of Uncharged IDPs"

### 1. Detailed steps of the analytical theory:

Consider the phase separated solution of FUS. It occupies volume V lattice cites (together with ions) while the rest of the solution in the dilute phase occupies the remaining volume $V_0$-V. The first region we call "inside" (in) and the polymer free region we call outside ("out"). We do not make distinction between $Na^+$ and $Cl^-$ ions (same interactions with polymers for simplicity). We will use Flory-Huggins approximation to present free energy of the system.

We consider a general two component phase separated system with volume of dense phase V, total volume $V_0$, total number of FUS chains, ions, and water molecules and the same in the dense phase correspondingly (we mark molecules in dense phase with symbol "in", molecules in dilute phase symbol "out" and total with symbol "0") $N_F^0,\ N_F^{In}, N_F^{In};\ N_I^0, N_I^{in}, N_I^{out}\ ; N_W^0\ , N_W^{in},\ N_W^{out}$. Free energy of the whole system is then:

$$F_{total} = V\left[ f\left( \frac{N_F^{in}}{V}, \frac{N_I^{in}}{V}, \frac{N_W^{in}}{V} \right) \right] + \left(V_0 - V\right)\left[ f\left( \frac{\left(N_F^0 - N_F^{in}\right)}{V_0 - V}, \frac{\left(N_I^0 - N_I^{in}\right)}{V_0 - V}, \frac{\left(N_W^0 - N_W^{in}\right)}{V_0 - V} \right) \right] \quad \text{(S1)}$$

Adding condition of incompressibility and assuming that monomers of each type are of the same volume $v_0$

$$v_0\left[N_F^{in}+N_I^{in}+N_W^{in}\right]=V$$
$$N_F^{in}+N_I^{in}+N_W^{in}=\frac{V}{v_0}$$
$$\rho_F^{in}+c_I^{in}+\rho_W^{in}=\frac{1}{v_0} \tag{S2}$$
$$\left(\rho_F^{in}v_0\right)+\left(c_I^{in}v_0\right)+\left(c_I^{in}v_0\right)=1$$
$$\phi_F^{in}+\phi_I^{in}+\phi_W^{in}=1$$
$$\phi_W^{in}=1-\phi_F^{in}+\phi_I^{in}$$

where $L$ is the polymerization index of FUS. We switched from densities to volume fractions using the relation $\phi=\rho v_0$ or $\phi=cv_0$. Equivalently we can derive the same equation for outside of the dense phase

$$\phi_W^{out}=1-\phi_F^{out}-\phi_I^{out} \tag{S3}$$

Equilibrium between dense and diluted phases of FUS calls for equal chemical potentials of FUS molecules between phases and equal chemical potentials of salt ions between the dense and diluted phases. In addition, the condition of equal osmotic pressure determines the partitioning between the phases, i.e. determines the volume of the dense phase V:

$$\frac{\partial F_{total}}{\partial N_I^{in}}=0 \;\rightarrow\; \mu_I^{in}\left(\phi_I^{in},\phi_F^{in}\right)=\mu_I^{out}\left(\phi_I^{out},\phi_F^{out}\right)$$
$$\frac{\partial F_{total}}{\partial N_F^{in}}=0 \;\rightarrow\; \mu_F^{in}\left(\phi_I^{in},\phi_F^{in}\right)=\mu_F^{out}\left(\phi_I^{out},\phi_F^{out}\right) \tag{S4}$$
$$\frac{\partial F_{total}}{\partial V}=0 \;\rightarrow\; \left(f\left(\phi_I^{in},\phi_F^{in}\right)-f\left(\phi_I^{out},\phi_F^{out}\right)\right)-$$
$$-\left(\phi_I^{in}\mu_I^{in}\left(\phi_I^{in},\phi_F^{in}\right)-\phi_I^{out}\mu_I^{out}\left(\phi_I^{out},\phi_F^{out}\right)\right)-\left(\phi_F^{in}\mu_F^{in}\left(\phi_I^{in},\phi_F^{in}\right)-\phi_F^{out}\mu_F^{out}\left(\phi_I^{out},\phi_F^{out}\right)\right)=0$$

The condition of conservation of total amounts of FUS and ions are:

$$V\rho_F^{in}+\left(V_0-V\right)\rho_F^{out}=N_F^0 \;\text{ or }\; \frac{V}{V_0}\rho_F^{in}+\left(1-\frac{V}{V_0}\right)\rho_F^{out}=\rho_F^0;$$
$$Vc_I^{in}+\left(V_0-V\right)c_I^{out}=c_I^0 \;\text{ or }\; \frac{V}{V_0}c_I^{in}+\left(1-\frac{V}{V_0}\right)c_I^{out}=c_F^0 \tag{S5}$$
$$\frac{V}{V_0}\varphi_{F,I}^{in}+\left(1-\frac{V}{V_0}\right)\phi_{F,I}^{out}=\phi_{F,I}^0$$

Where $\rho_F^0$ and $c_I^0$ are initial total concentration of FUS and ions respectively.

Excluding water using the incompressibility conditions leaves us with effective free energy in the Voorn-Overbeek approximation[1] (**Eq. (2)** of the Main text) which treats polymers in the FH approximation and ion free energy in the Debye-Huckel approximation. As pointed out in the main text $\tilde{\chi}$ in Eq.2 are effective interactions between FUS and Ions – effective because water interactions are excluded through incompressibility conditions.

$$\begin{aligned}\tilde{\chi}_{FF} &= \chi_{FF} + \chi_{WW} - \chi_{WI} - \chi_{WF} - T \\ \tilde{\chi}_{FI} &= \chi_{FI} - \chi_{IW} - \chi_{FW} + 2\chi_{WW} - T\end{aligned} \tag{S6}$$

$F_{DH}$ is free energy of ion-ion interactions in the Debye-Huckel approximation which will be detailed later.

In the regime where both FUS and ion concentrations are low $\phi_I{}^{in} + \phi_F{}^{in} \ll 1$ we can omit water entropy term in **Eq. (2)** of Main Text. Denoting $x = \frac{\phi_I^{in}}{\phi_I^{out}} = \frac{c_I^{in}}{c_I^{out}}$ and substituting into **Eq. (S6)** we get for the chemical equilibrium for the ions (see **Eq. 3** of main text and accompanying discussion)

$$\begin{aligned}T\ln x - A\sqrt{c_I^0}\left(\sqrt{x}-1\right) + \tilde{\chi}_{Fi}\phi_F^{in} &= 0 \\ T\ln x - A\sqrt{c_I^0}\left(\sqrt{x}-1\right) &= -\tilde{\chi}_{Fi}\phi_F^{in}\end{aligned} \tag{S7}$$

Maxima of the curves shown in **Figure S1(a)** are reached at

$$\frac{\phi_I^{in}}{\phi_I^{out}} = x_{\max} = \frac{4T^2}{A^2 c_0} = \frac{Q^2}{c_0} \text{ where } Q = \frac{2T}{A} \tag{S8}$$

Maximal value of $\phi_F^{\max}$ possible is given by:

$$\begin{aligned}&T\ln x_{\max} - A\sqrt{c_0}\left(\sqrt{x_{\max}}-1\right) = -\tilde{\chi}_{FI}\phi_F^{max} \\ &\phi_F^{max} = -\frac{T\ln x_{\max} - A\sqrt{c_0}\left(\sqrt{x_{\max}}-1\right)}{\tilde{\chi}_{FI}} = -\frac{T\ln\frac{4T^2}{A^2c_0} - \frac{4T^2}{A\sqrt{c_0}} + A\sqrt{c_0}}{\tilde{\chi}_{FI}} = \\ &-\frac{\ln\frac{Q^2}{c_0} - \frac{2Q}{\sqrt{c_0}} + \frac{2}{Q}\sqrt{c_0}}{\frac{\tilde{\chi}_{FI}}{T}}\end{aligned} \tag{S9}$$

The plot of $\phi_F^{max}$ vs salt concentration is shown in **Figure S1(b)**

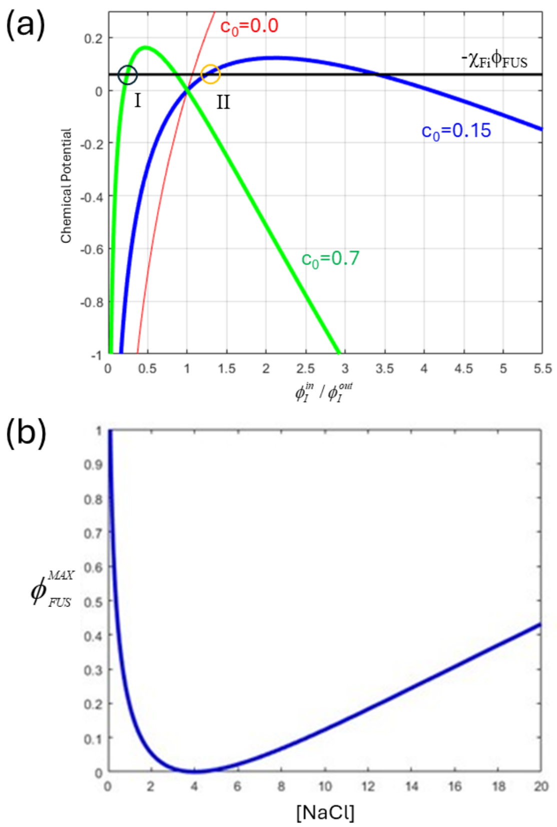


**Figure S1. (a) Graphical solutions of Eq. (S7):** The red line: no salt charges, (just inert crowders instead of charged ions), the blue line $\phi_I^{out} = 0.15$ ,the green line $\phi_I^{out} = 0.7$, the black line constant RHS of $-\chi_{Fi}\phi_{FUS}$. The intersections marked with 'I' and 'II' are the stable solutions. **(b) Maximal density (volume fraction) of FUS at which Eq.(S9) has nontrivial solutions** as a function of [NaCl] concentration (in arbitrary units). At higher volume fractions of FUS equilibrium is possible only at $\phi_I^{in} / \phi_I^{out} = 1$ and $\phi_{FUS} = 0$ which is the case at intermediate salt concentrations where non-trivial equilibrium is achieved at very low FUS volume fractions indistinguishable from random coil leading to de-collapse upon increase of [NaCl] and collapse again at much higher [NaCl].

It is clear from this plot that at low salt concentration the chain can collapse (provided that free energy of the collapsed state is lower than random coil, see below), at intermediate concentrations it can be only random coli and at higher concentrations the chain can collapse again as MAX density of FUS again rises. That explains, in principle two reentrant transitions. At $c_0 < Q$ maximum is at $x = \phi_I^{in} / \phi_I^{out} > 1$ .

The plots on Figure S1(a) show that for the purpose of solving the equation in the relevant range of x we can approximate the chem potential function as a parabola:

$$-\left(x-1\right)(x-x^{*})+\chi_{Fi}\phi_F^{\max}=0 \text{ where } x^{*}=2x_{\max}-1=\frac{2Q^2}{c_0}-1 \quad \text{(S10)}$$

Solving for x we get:

$$x=x_{\max}-\sqrt{\left(x_{\max}-1\right)^2+\chi_{Fi}\phi_F^{\max}} \quad \text{(S11)}$$

Giving us the result for concentration of ions inside the FUS polymer as

$$\phi_I^{in}=\phi_I^{out}\left(\frac{4T^2}{A^2\phi_I^{out}}-\sqrt{\left(\frac{4T^2}{A^2\phi_I^{out}}-1\right)^2+\tilde{\chi}_{Fi}\phi_F^{\max}}\right) \quad \text{(S12)}$$

Now consider the onset of collapse i.e. $\phi_F^{in}\ll 1$. In this case we expand the term with the square-root, in Eq.(S15) and get finally:

$$\phi_I^{in}=\phi_I^{out}-\frac{\chi_{Fi}\left(\phi_I^{out}\right)^2\phi_F^{in}}{\frac{4T^2}{A^2}-\phi_I^{out}}\approx\frac{A^2\chi_{Fi}\left(\phi_I^{out}\right)\phi_F^{in}}{4T^2} \text{ when } \phi_I^{out}\leq 1 \quad \text{(S13)}$$

Substituting this expression for $\phi_{out}$ into the original FH **Eq. (2)** of main text we get

$$F=M_1\left[\frac{T}{N}\phi_F^{in}\ln\phi_F^{in}+\left(\chi_{FF}-\frac{A^2\chi_{Fi}{}^2(\phi_I^{out})^2}{4T^2}\right)\left(\phi_F^{in}\right)^2+C_3\left(\phi_F^{in}\right)^3\right] \quad \text{(S14)}$$

Here we see that expulsion of ions effectively leads to additional attraction between FUS monomers by renormalizing its second virial coefficient of interactions

$$\tilde{\chi}_{FF}^{eff}(\phi_I^{out})=\left(\chi_{FF}-\frac{A^2\chi_{Fi}{}^2(\phi_I^{out})^2}{4T^2}\right) \quad \text{(S15)}$$

When

$$\chi_{FF}^{eff}(\phi_I^{out})=\left(\chi_{FF}-\frac{A^2\tilde{\chi}_{FI}{}^2(\phi_I^{out})^2}{4T^2}\right)=0 \text{ i.e when}$$
$$\phi_I^{out}=\frac{2T}{A}\frac{\sqrt{\tilde{\chi}_{FF}}}{\tilde{\chi}_{FI}} \quad \text{(S16)}$$

The chain collapses, i.e. it reaches an effective $\theta$-point.

### 2. Sequence Analysis:

**Table S1(a). Fraction of positively charged residues (f+), fraction of negatively charged residues (f-), total charge (Net Q), and net charge per residue (NCPR) for the five full length constructs used in the experimental study by Krainer *et al.*[2]**

| Systems | Residues | f+ | f- | Net Q | NCPR |
|---|---|---|---|---|---|
| FUS | 1-526 | 0.097 | 0.070 | +14 | +0.026 |
| TDP-43 | 1-414 | 0.097 | 0.106 | -4 | -0.0096 |
| Brd4 | 1-1362 | 0.128 | 0.110 | +25 | +0.0183 |
| Sox2 | 1-317 | 0.107 | 0.066 | +13 | +0.041 |
| A11 | 1-505 | 0.101 | 0.099 | +1 | +0.002 |

**Table S1(b). The same parameters calculated for the disordered (low complexity) droplet promoting regions of the five systems listed in Table S1(a).**

| Systems | Residues | f+ | f- | Net Q | NCPR |
|---|---|---|---|---|---|
| FUS | 1-163 | 0.0 | 0.012 | -2 | -0.012 |
| | 164-286 | 0.098 | 0.081 | +2 | +0.016 |
| TDP-43 | 261-303 | 0.116 | 0.047 | +3 | +0.071 |
| | 341-373 | 0.030 | 0.030 | 0 | 0.000 |
| Brd4 | 1-58 | 0.086 | 0.069 | +1 | +0.017 |
| | 174-229 | 0.107 | 0.018 | +5 | +0.091 |
| | 242-352 | 0.144 | 0.081 | +7 | +0.063 |
| | 463-615 | 0.216 | 0.196 | +3 | +0.019 |
| Sox2 | 1-43 | 0.116 | 0.070 | +2 | +0.046 |
| | 243-266 | 0.083 | 0.042 | +1 | +0.043 |
| | 297-317 | 0 | 0 | 0 | 0.000 |
| A11 | 1-38 | 0 | 0 | 0 | 0.000 |
| | 84-199 | 0.017 | 0.009 | +1 | +0.009 |

### 3. Pairwise Interaction Energetics Figures:

In this section, we plot the pairwise interaction energetics of different pairs namely, inter-protein ($E_{FUS}$), protein-water ($E_{FUS-W}$), protein-ion ($E_{FUS-Na^+}$ or $E_{FUS-Cl^-}$), and inter-ion interactions ($E_{ION-ION}$); for both the original system (**Figure S2**) and a control system where the ionic charges are removed (**Figure S3**). For the latter, there is no distinction between sodium and chloride ions. Hence, instead of $E_{FUS-Na^+}$ or $E_{FUS-Cl^-}$ we use $E_{FUS-ION}$. All the energy values are a combination of electrostatic (Coulomb) and van der Waals (Lennard-Jones) interactions, and are normalized with respect to the number of FUS chains, *N*.

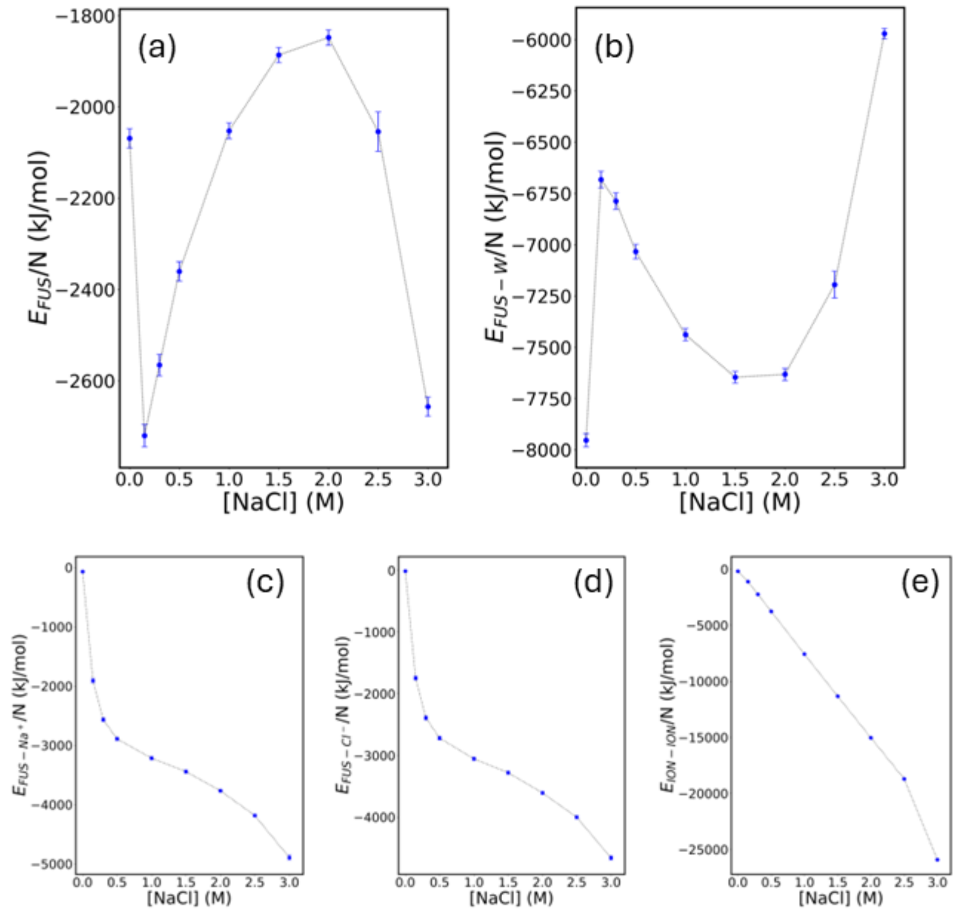


**Figure S2. Pairwise interaction energetics against the solution's ionic strength (when the ions carry their natural charges):** (a) Inter-protein interaction sharply drops at 0.15 M [NaCl] followed by a gradual increase, reaching a maximum, and again dropping sharply at 3 M [NaCl]. The sharp decreases indicate condensation. (b) The protein-water interaction energy shows a sharp increase at 0.15 M [NaCl] followed by a gradual decrease, reaching a minimum, and again increasing at 3 M [NaCl]. (c) Protein-sodium ion and (d) protein-chloride ion interaction energetics exhibit monotonic decrease with the increasing salt concentration. (e) Inter-ion interactions also show a monotonic decrease with respect to solution's ionic strength.

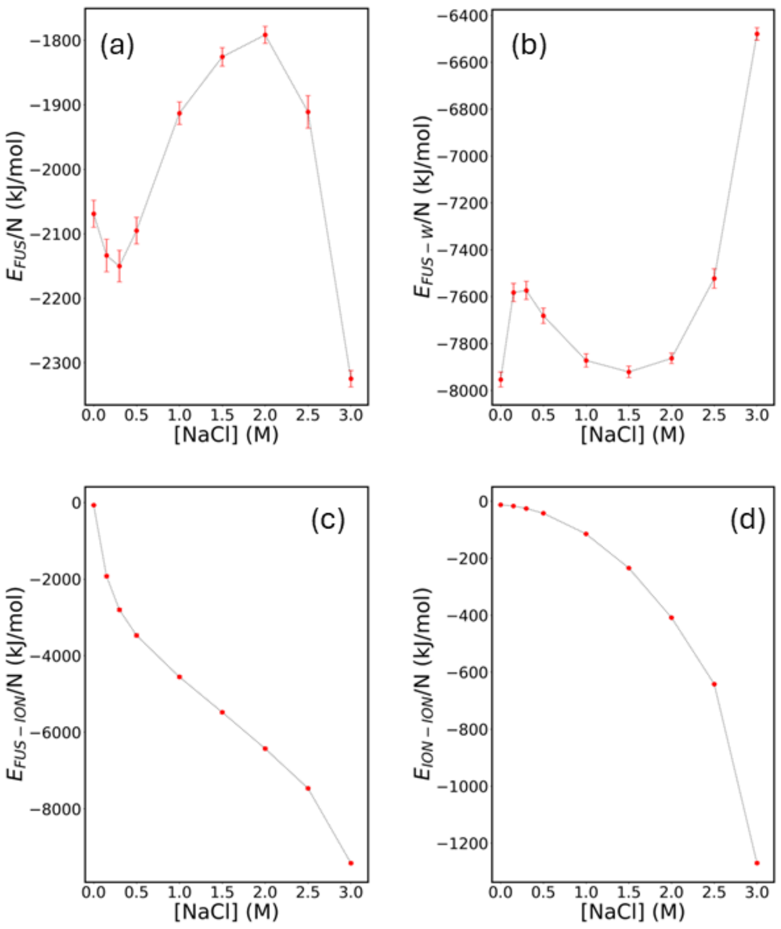


**Figure S3. Pairwise interaction energetics against the solution's ionic strength (when electrostatic interactions are 'turned off' that is, ionic charges are set to '0'):** (a) Inter-protein interaction shows a little dip around 0.15 M [NaCl] before increasing, reaching a maximum, and decreasing sharply at 3 M [NaCl]. The initial dip indicates enthalpic stabilization to some extent but not condensation. (b) Protein-water interaction initially shows a slight increase followed by gradual decrease, reaching a minimum, and a sharp increase at 3 M [NaCl]. (c) Protein-ion interaction and (d) inter-ionic interaction show monotonic decrease with increasing [NaCl].

4. **Condensation in System-2 Starting from a Dispersed State:**

System-2, where the FUS charges are removed but salt ion charges are kept intact, exhibit LLPS. In the main text (**Figure 3b**) we showed the final state of the system starting with a preformed droplet state. To rule out a possible kinetic trap we carried out additional simulation starting with a fully dispersed state and observed spontaneous LLPS within a few microseconds of coarse-grained timescale (**Figure S4**).

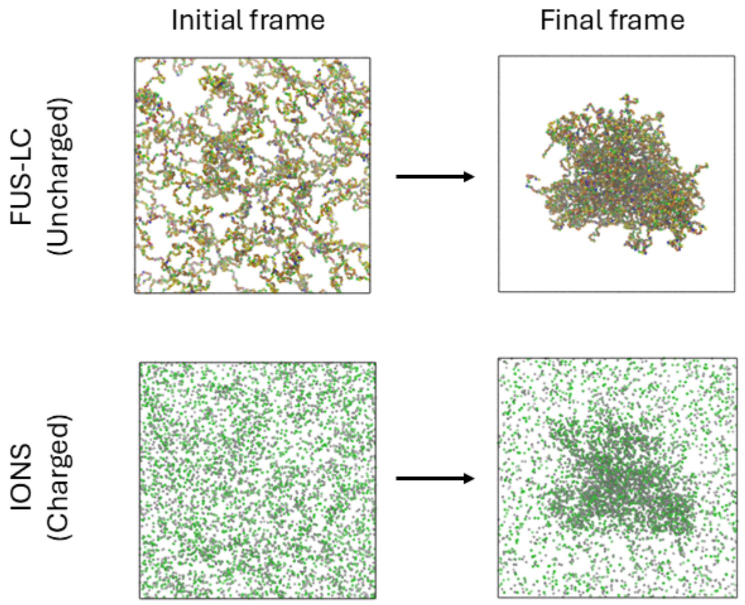


**Figure S4. LLPS of uncharged FUS-LC with charged 0.15 M NaCl, starting from a fully dispersed state:** Top panel shows the distribution of FUS chains at their initial (dispersed) and final (condensed) state. The bottom panel shows the distribution of Na+ (grey) and Cl- (green) ions for the same timeframes as above.

5. **Comparison between MARTINI-3 and Madrid-2019 model contact ion pairs:**

We have simulated NaCl ions in water at two different high salt concentrations, namely 1.50 M, and 3.00 M. At T=298K and p=1 bar, from classical MD simulations, we have obtained contact ion pair ($n_{CIP}$) from the partial radial distribution function, g(r) between Na+ and Cl- as follows:

$$n_{CIP} = 4\pi\rho \int_0^{r_{cut}} dr\, r^2 g(r) \tag{S17}$$

where $r_{cut}$ is the cut-off distance up to which we integrate g(r) and $\rho$ is the number density of either $Na^+$ or $Cl^-$. We compare the MARTINI-3 results with one of the best available forcefields for ionic solution (Madrid-2019 with TIP4P/2005 water model)[3] that predicts NaCl solubility close to 6 M. For comparison purposes, we set $r_{cut}$=0.6 nm which is close to a minimum in g(r). The results are tabulated below:

**Table S2. Contact ions pairs obtained from MARTINI-3 (coarse-grained) and Madrid (atomistic) force fields for two different high ionic strengths.**

| MARTINI-3 results | | |
|---|---|---|
| Systems | ρ ($nm^{-3}$) | $n_{CIP}$ |
| 1.50 M | 0.93 | 0.95 |
| 3.00 M | 1.85 | 1.49 |
| Madrid results | | |
| Systems | ρ ($nm^{-3}$) | $n_{CIP}$ |
| 1.50 M | 0.90 | 1.04 |
| 3.00 M | 1.80 | 1.95 |

From **Table S2**, we notice that the number densities (after NpT simulation) obtained from MARTINI force-field, on average, are only 3% higher than those obtained from the atomistic force field. For the MARTINI model, the values of $n_{CIP}$ are even lower than those of Madrid model. A higher value of $n_{CIP}$ denotes an increased 'crowding' of cations (or anions) around anions (or cations). We provide the radial distribution plots below (**Figure S5**), where the solid lines denote Atomistic force field and dashed lines denote coarse-grained force field.

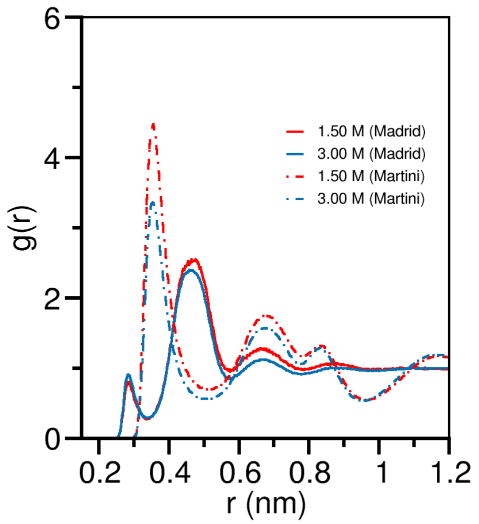


**Figure S5. Radial distribution function (RDF) comparison between atomistic and MARTINI models:** RDF of Cl- ions with respect to Na+ ions in two different salt concentrations, by using Martini (coarse-grained) and Madrid (atomistic) models.

The two peaked fine structures of g(r) in the atomistic force field (centered around 0.28 nm and 0.47 nm) are merged to one peak structure (centered around 0.36 nm). This can be attributed to the difference in the van der Waals radii ($\sigma$) of ion and water beads. For example, in the atomistic force field $\sigma_{Na^+} = 0.2217$ nm and $\sigma_{H_2O} = 0.3159$ nm; whereas in the MARTINI coarse-grained description $\sigma_{Na^+} = 0.354$ nm and $\sigma_{H_2O} = 0.470$ nm. As a result, the finer peak structures in g(r) disappear and a 'coarser' peak appears in between. Nevertheless, the value of $n_{CIP}$ remains comparable between the two force fields.

### 6. LLPS of full-length FUS:

In the main text, we discussed the reentrant LLPS of the low complexity domain of FUS. Here, we provide the LLPS propensity of full-length FUS at three different salt concentrations, namely, 0.15 M, 1.5 M, and 3.0 M. We find that, the full-length FUS exhibits condensation at 0.15 M and 3.00 M, but not at 1.5 M. This corroborates well with the observation with the LLPS of FUS-LC only. However, the phase boundaries may change/shift if one compares the phase behavior of the full-length FUS and the LC region. The comparison can only be in a qualitative sense.

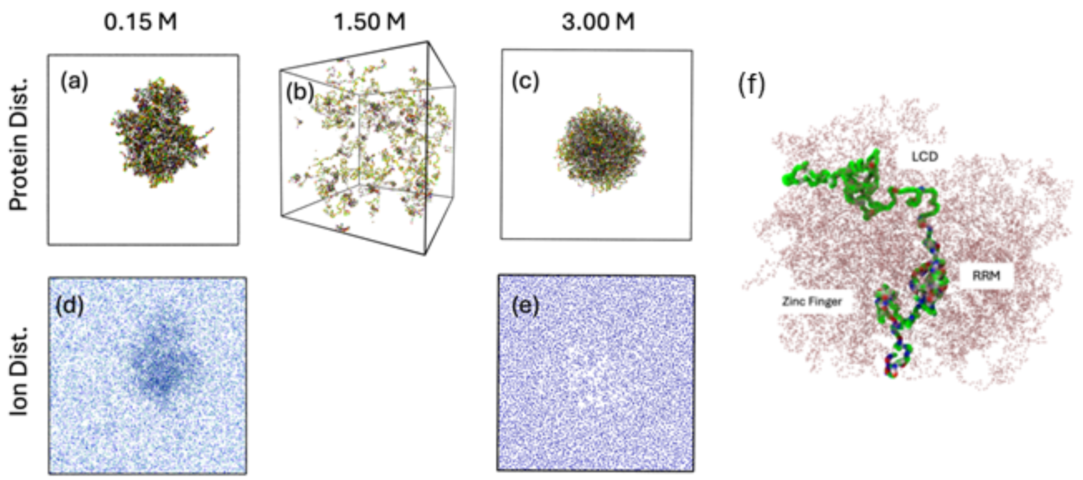


**Figure S6. MD simulation of Full-length FUS:** (a), (b), and (c) are representative snapshots of the system after 3 [illegible]s MD simulation for three different salt concentrations: 0.15 M, 1.5 M, and 3.0 M respectively. (d) and (e) are the spatial distribution of ions inside the box at the exact same time-point as the condensate snapshots. The observations by simulation of the FUS-LC domain are valid when full length FUS is used. (f) Magnified view of the condensate in panel 'a' where the conformation of a single full-length FUS is highlighted.

### 7. LLPS of FUS-LCD at less than 0.15 M NaCl

To comment on the threshold salt concentration of the LLPS of FUS-LCD, we have carried out additional coarse-grained simulations at $[C_{ion}]$ = 0.05 M and 0.10 M for 5 µs starting form a completely dispersed state. We observe LLPS at $[C_{ion}]$ = 0.10 M but not at $[C_{ion}]$ = 0.05 M (**Figure S7**). This suggests that the threshold lies in between 0.05 M < $[C_{ion}^{threshold}]$ < 0.10 M. We reiterate that these results are with FUS-LCD and the threshold might differ for full length FUS. In **Figure S7(a)** we show the time evolution of the radius of gyration of the entire protein system and in **Figure S7(b)** we show the distribution of number of clusters.

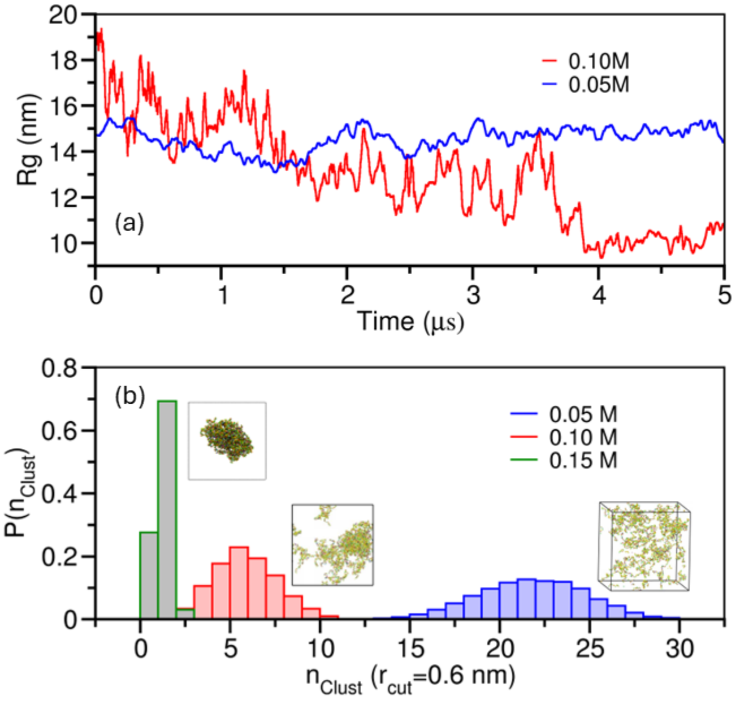


**Figure S7. LLPS propensity of FUS-LCD at salt concentrations lower than 0.15 M:** (a) Time evolution of the radius of gyration (Rg) of all the proteins. At 0.05 M the Rg shows no sign of decrease along the trajectory whereas at 0.10 M the Rg significantly drops around 4 ms. (b) The distribution of numbers of clusters in the system shows right shifted distribution at 0.05 M indicating the presence of several smaller clusters. At 0.10 M the distribution shifts to the left indicating the onset of LLPS (snapshot beside the distribution) and as a reference, the sharply peaked distribution is at 0.15 M.

### 8. Validation from atomistic simulations:

we have performed atomistic simulations with CHARMM36 forcefield and TIP3P water model. We have utilized the back-mapping algorithm to convert the equilibrated coarse-grained condensates into atomistic resolution (for details please see https://cgmartini.nl/docs/tutorials/Martini3/Backward/). We have additionally modified the Lennard-Jones parameters of $Na^+$ and $Cl^-$ ions following the work of Yagasaki *et al.*[4] who obtained a solubility of 6.1 ± 0.30 mol/kg for NaCl in TIP3P water model.

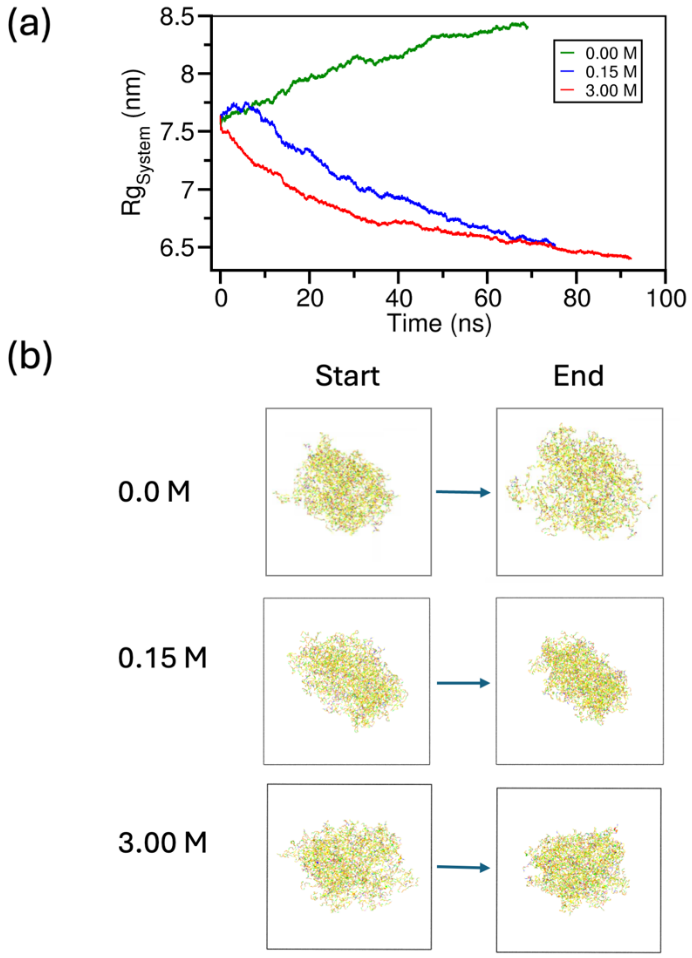


**Figure S8. Atomistic simulation of a preformed droplet at three salt concentrations.** (a) Time evolution of Rg of the protein system during atomistic MD simulation shows a gradual increase in Rg for 0.0 M salt and a gradual decrease in Rg indicating compaction of the condensate for 0.15 M and 3.0 M, compared to its backmapped configuration from MARTINI. (b) Snapshots of the systems before and after atomistic MD simulations at three different salt concentrations showing swelling for 0.0 M and compaction for 0.15 M and 3.0 M during the atomistic simulation.

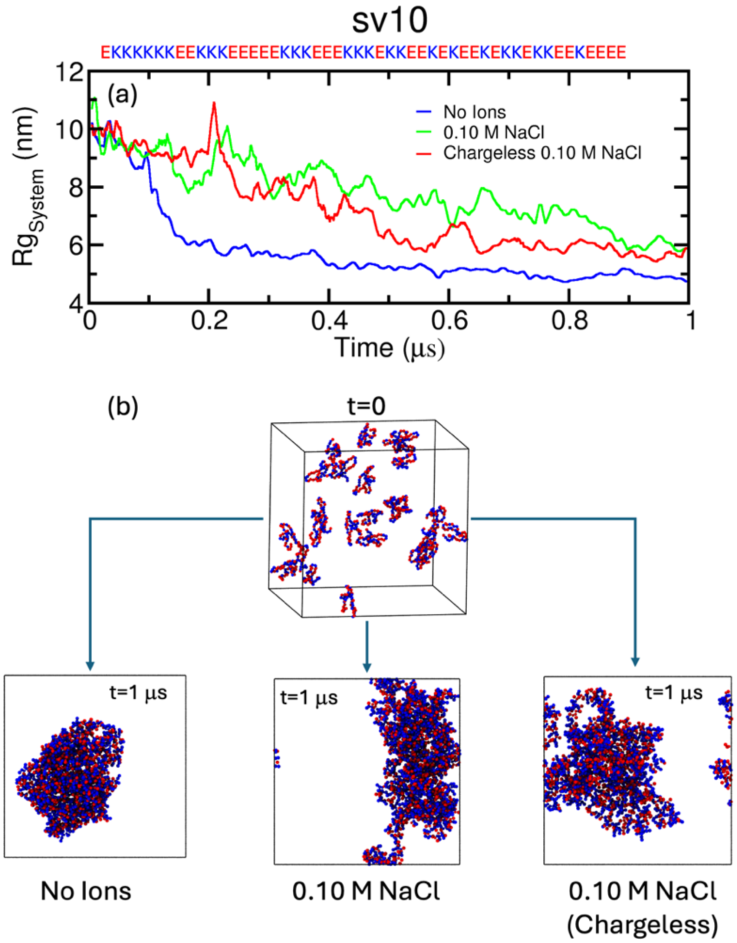


**Figure S9. Phase separation of 'sv10' polyampholyte:** (a) Time evolution of Rg of the polyampholyte system from 0 to 1 μs. For the system with no ions, Rg shows a sharp decrease around 150 ns indicating fast self-condensation without the help of external ions. For the 0.10 M charged/uncharged NaCl system the decrease in Rg is more gradual. (b) Representative snapshots of the sv10 polyampholyte system at t=0 and t=1 μs for three systems. The condensate is more compact when no ions were present, compared to 0.10 M NaCl charged/uncharged systems.

3. Since polyampholytes can form condensates at 0 M salt, the re-entrant phase diagrams are different than FUS-LC type 'nearly uncharged' sequences. A simple schematic comparison is drawn in **Figure S10**. Panel (a) shows a representative salt concentration dependent phase diagram for FUS-LC type 'nearly uncharged' IDP sequences where there are 4 distinct regions, out of which $\alpha, \gamma$ are 'No LLPS' zones;

and $\beta, \delta$ are the 'LLPS' zone. Along the [salt] axis, the system can exhibit three transitions as shown by purple arrows, making it a double-reentrant phase transition.

On the other hand, panel (b) shows a typical re-entrant phase diagram for polyampholyte sequences, that are charged but with a net neutrality. Here one finds three distinct regions: $\beta', \delta'$ are the 'LLPS' zones and $\gamma'$ is the 'No LLPS' zone. The system can make two transitions along the [salt] axis, making it a single-reentrant phase diagram.

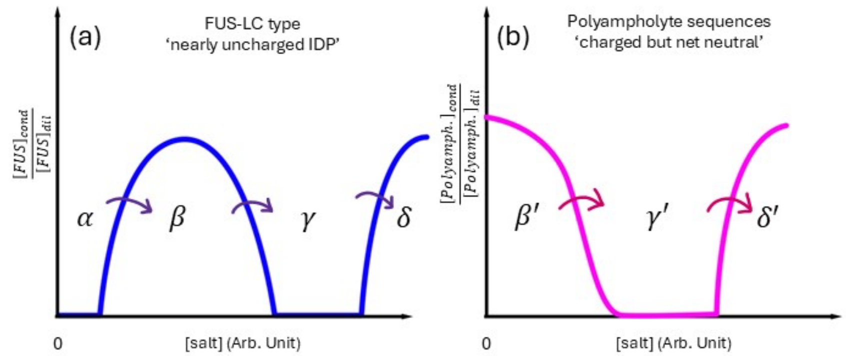


**Figure S10. Schematic phase diagrams of (a) FUS-type nearly uncharged IDPs and (b) polyampholyte systems, against ionic strength.**
